# VIP-OT: Dissecting Single-Cell Biochemical State Dynamics under Perturbation via Vibrational Painting and Optimal Transport

**DOI:** 10.64898/2025.12.09.693255

**Authors:** Xuemeng Li, Stefan G. Stark, Xinwen Liu, Mian Wei, Zhilun Zhao, Jianhuan Qi, Xingjian Chen, Thai Nam Son Dang, Yinghan Wu, Ke Zhang, Wei Min, Jian Shu

## Abstract

Dissecting the heterogeneous response of individual cells towards genetic and chemical perturbations is central to understanding the dynamic functions of cells and multicellular systems. However, characterizing and modeling how individual cells transition between different states remains a major challenge. Vibrational imaging provides high-content, biochemically informative, label-free molecular fingerprints of single cells but it remains at its infancy for dynamic or predictive analysis of cell state transition. Here, we introduce Vibrational Painting-Optimal Transport (VIP-OT), an integrated experimental-computational framework that overcomes this fundamental limitation. VIP-OT couples multiplexed infrared (IR) and Raman imaging with optimal transport to computationally reconstruct single-cell perturbation trajectories from unpaired population snapshots. When applying to over 22,000 single-cell spectra profiles from human breast adenocarcinoma cells under 16 drug treatments, this framework can retrospectively trace drug response heterogeneity back to baseline metabolic states. We leverage the inferred cell pairings to develop a machine learning model that accurately predicts the full post-treatment metabolic state of individual cells from their pre-treatment spectra. Furthermore, by modeling transitions across dose gradients, we introduce the concept of Spectral Velocity to map dynamic response trajectories and resolve drug combination effects into distinct, path-dependent molecular routes. Together, VIP-OT opens a new direction for dissecting heterogeneous perturbation responses at single cell resolution and serves as a foundation for building virtual simulators of cells under perturbations through high-throughput, high-content, and low-cost vibrational imaging, as well as interpretable *in silico* modeling.

## Main

Understanding how individual cells respond to genetic or chemical perturbations lies at the heart of every aspect of biology, ranging from uncovering molecular functions of multicellular systems to the drug discovery pipeline^1,2^. However, such responses are often heterogeneous due to a range of intrinsic and extrinsic factors, including variations in metabolic or transcriptional states, local microenvironments, and nonlinear gene-gene interactions^3,4^. Capturing and modeling this heterogeneity is essential for identifying responsive subpopulations and constructing predictive frameworks of cell behavior. For example, drug-resistant clones can emerge from minor subpopulations in cancer^5^, rare immune subsets can disproportionately drive host defense^6^, and stem cell populations display variable differentiation propensities even under uniform conditions^7,8^. Single-cell transcriptomic and imaging studies have repeatedly shown that such heterogeneity is often masked in bulk measurements, yet proves decisive for treatment outcomes and developmental trajectories^9,10^. In this context, the concept of the AI Virtual Cell (AIVC) has recently emerged a computational simulator capable of modeling and forecasting cellular behaviors and functions across modalities and scales^11^. Realizing this vision, however, demands massively scalable techniques capable of capturing perturbation responses at single-cell resolution, together with analytical methods that can robustly address their high-dimensional, continuous, and heterogeneous nature^2,11–13^.

A range of single-cell technologies have been developed to characterize response heterogeneity. For example, single-cell sequencing offers an unbiased view of cellular diversity and enables detection of rare subpopulations but suffers from high-cost and batch effects^14,15^. Mass spectrometry-based methods provide high-content molecular profiling, yet approaches such as CyTOF rely heavily on antibody specificity, and involve labor-intensive, low-throughput workflows with high costs^16^. Fluorescence-based imaging provides high sensitivity and subcellular spatial resolution, but is limited to a small number of predefined targets, and fluorescent probes may perturb native cellular physiology^17,18^.

Vibrational spectroscopic imaging offers an alternative approach by leveraging endogenous biomolecules as intrinsic signals, enabling functional analysis of single cells. The unique “fingerprint” spectra generated by quantized vibrations of chemical bonds allow identification of molecular species within each cell, typically measured using infrared (IR) or Raman spectroscopy^19^. By coupled with vibrational metabolic probes, the sensitivity and specificity of this approach can be markedly improved^20–24^. Building on this principle, we recently developed VIBRANT, a vibrational imaging framework that integrates multiplexed metabolic probes with single-cell IR imaging to capture phenotypic responses to chemical perturbations^25^. It accurately classified drug Mechanisms of action (MoA) from single-cell spectra with robust cross-batch performance, and also demonstrated discovering drug candidates with novel MoAs. VIBRANT enables high-throughput and high-content profiling of metabolically labeled single cells, offering a scalable and cost-effective platform with minimal batch effects. As such, VIBRANT addresses several key limitations faced by current single-cell technologies, providing a new direction for single-cell metabolic analysis.

However, while vibrational spectroscopic imaging is in principle nondestructive and compatible with live-cell measurements, achieving high-throughput longitudinal tracking across thousands of perturbed single cells remains technically prohibitive. Continuous live-cell imaging at this scale is challenged by phototoxicity^26–28^, limited acquisition speed^19,20,29^, and difficulties in maintaining cell stability and registration over long periods^30,31^. For this reason, VIBRANT relies on fixed-cell measurements, where each experiment yields an independent population snapshot without natural pairing between control and treated cells, precluding longitudinal tracking of the same cell before and after perturbation. Consequently, it remains impossible to directly model how an individual cell transitions between states, constraining researchers from reconstructing dynamic state transition trajectories or predicting future perturbation outcomes at single-cell resolution.

To overcome this barrier, we propose reframing the challenge from a distributional perspective. Instead of tracking individual cells, which is technically prohibitive, we can consider each cell population as an empirical probability distribution in the high-dimensional space of its spectral profiles. The problem then becomes how to mathematically model the transition between these two distributions to infer cell-level responses. For this purpose, we turned to optimal transport (OT), a mathematical framework originating from Monge’s 18th-century problem of efficient mass relocation and later formalized by Kantorovich^32^. Recently, OT has proven transformative in single-cell biology: we first introduced OT for single-cell genomics applications and developed Waddington-OT to reconstruct developmental trajectories from single-cell sequencing^33^; Cell-OT predicted genetic and chemical perturbation outcomes^34^; and others have developed related approaches to map cell states across time and space or dissected communication in spatial transcriptomics^35–38^. These studies establish OT as a powerful tool for aligning heterogeneous cell populations and inferring transitions^39–41^, but their scope has been confined to single-cell omics, leaving its potential for vibrational imaging unexplored.

Building on this framework, we specifically adapted OT to the context of vibrational imaging. To our knowledge, this represents the first application of OT to vibrational imaging. Rather than imposing one-to-one correspondences, we formulated perturbation-induced changes as a distributional alignment problem: OT identifies the probabilistic coupling between control and treated populations that minimizes the overall *spectral* displacement. This pseudo-pairing framework preserves global population structure while enabling per-cell inference without requiring paired measurements.

Here, we develop an integrated experimental and computational framework, named, Vibrational Painting-Optimal Transport (VIP-OT) through multiplexed vibrational imaging coupled with OT-based *in silico* modeling. Applying VIP-OT to over 22,000 single-cell spectral profiles acquired via IR and Raman imaging from human breast adenocarcinoma cells under 16 drug treatments, we uncover how phenotypic heterogeneity in drug response is shaped by baseline metabolic features, enabling retrospective tracing of subpopulation-specific sensitivities. By leveraging OT-derived pairings, we further predict the metabolic response of individual cells under specific perturbations. Moreover, by modeling dose-dependent transitions, VIP-OT reconstructs spectral response trajectories and introduces the concept of *Spectral Velocity* (SVL) to quantify the magnitude and direction of cellular state changes. Finally, by resolving drug combination paths, VIP-OT reveals distinct drug mechanisms at the single-cell level.

Together, VIP-OT transforms static vibrational imaging into a dynamic and predictive framework for single-cell perturbation analysis. VIP-OT not only overcomes the fundamental pairing barrier in vibrational data but also establishes a generalizable strategy for reconstructing and predicting cellular responses. This approach opens new avenues for building virtual simulators to understand the biochemical activities and functions of physical cells.

## Result

### Inferring single-cell perturbation responses on high dimensional vibrational profiles

To infer single-cell perturbation responses from static vibrational snapshots, we developed Vibrational Painting-Optimal Transport (VIP-OT). The experimental arm of VIP-OT enhances biochemical specificity by incorporating three IR- and Raman-active metabolic probes—¹³C-amino acids, azido-palmitic acid, and deuterated oleic acid—to simultaneously report on protein synthesis, saturated fatty acid metabolism, and unsaturated fatty acid metabolism, respectively^22,25^. The workflow (**Fig. 1**) begins with culturing human breast cancer cells (MDA-MB-231 cells) with these probes while subjecting them to drug perturbations. Following treatment, cells are fixed, and their molecular composition is captured using either FTIR or spontaneous Raman spectroscopy (Methods). Single-cell segmentation and spectral extraction yield a population of high-dimensional vibrational fingerprints and profiles for each condition, representing the biochemical composition and metabolic state of each individual cell. Crucially, these population snapshots lack inherent pairings between control and treated cells. To overcome this challenge, VIP-OT models each cell population as an empirical distribution in a shared high-dimensional spectral space ***χ***. The control and treated populations, donated *µ* and *v*, represent ensembles of single-cell spectra sampled from their respective conditions. We then apply Optimal transport (OT) to seek a mapping T from control states to treated states by minimizing total spectral displacement^42,43^. Rather than enforcing a deterministic pairing, OT computes a soft probabilistic coupling *γ* that defines the most likely correspondence between the two populations under a principle of minimal effort (Methods).

**Fig. 1.**
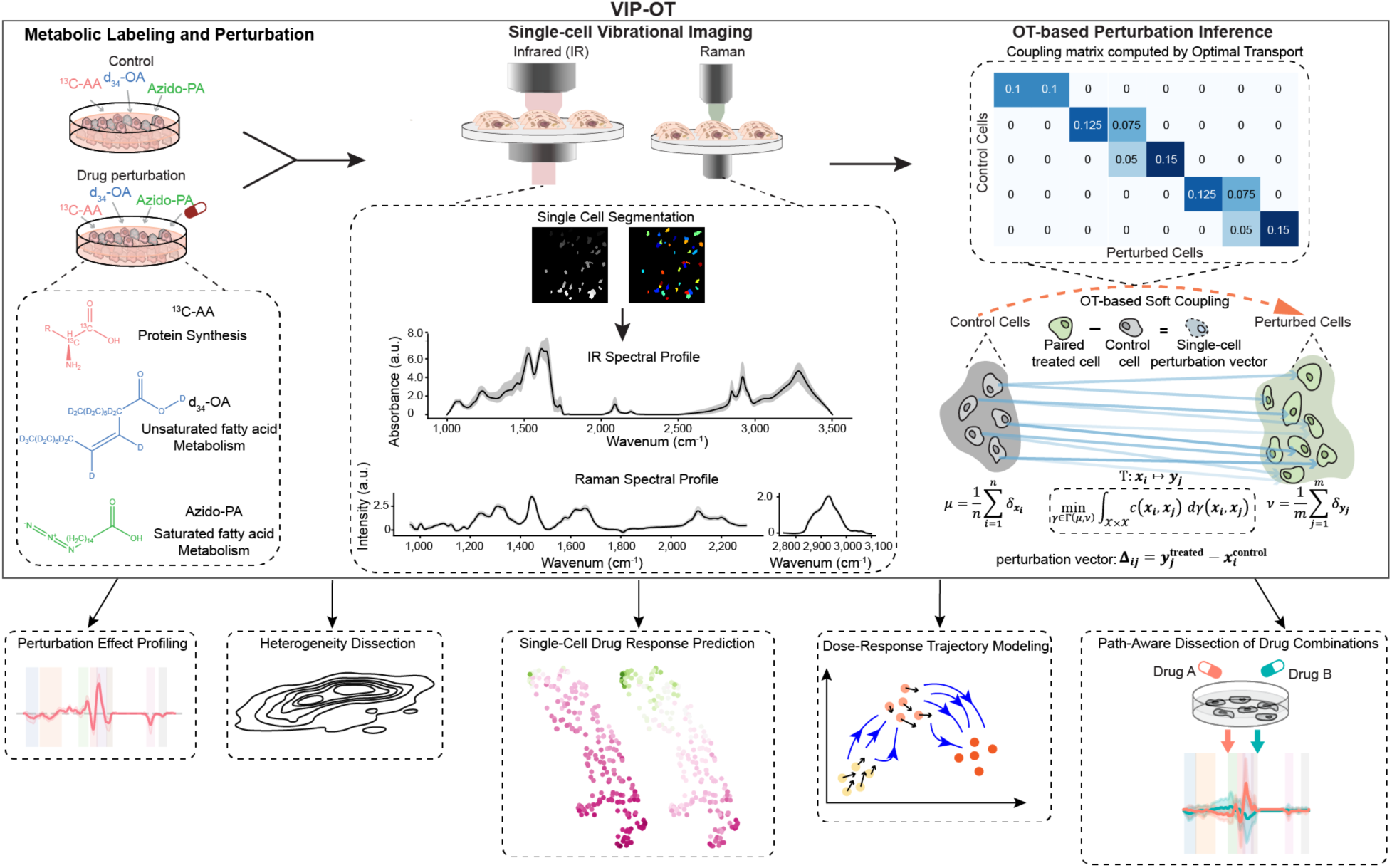
VIP-OT enables single-cell perturbation inference through optimal transport. Overview of the VIP-OT workflow. VIP-OT integrates metabolic labeling, single-cell vibrational imaging, and optimal transport to quantify drug-induced metabolic changes at single-cell resolution. MDA-MB-231 cells were labeled with three vibrational probes—¹³C-amino acids, azido-palmitic acid, and deuterated oleic acid—to monitor protein synthesis, saturated fatty acid metabolism, and unsaturated fatty acid metabolism, respectively. After drug perturbation, cells were fixed and imaged using either Fourier-transform infrared (FTIR) or spontaneous Raman spectroscopy, and single-cell spectra were extracted (Methods). To infer perturbation effects without paired measurements, OT was used to compute a soft coupling between control and treated populations, identifying minimal-cost spectral mappings. This coupling serves as the computational backbone for a suite of downstream analyses including drug effect profiling, heterogeneity dissection, single cell drug response prediction, dose–response trajectory modeling, and path-aware combination analysis.

From the resulting coupling, we derived perturbation vectors *Δ* for each control cell, expressed as the displacement toward its most likely treated counterparts and weighted by *γ* (Methods). These perturbation vectors form the computational backbone of VIP-OT. By enabling direct inference of cell-specific perturbation vectors from unpaired snapshots, VIP-OT overcomes a fundamental limitation of fixed-cell vibrational imaging: making it possible, for the first time, to reconstruct single-cell state transitions, trace baseline-dependent heterogeneity, predict individualized drug responses, and resolve continuous trajectories of dose and combination effects, as illustrated in **Fig. 1**.

### VIP-OT–inferred perturbation vectors encode biologically relevant mechanism of action

To evaluate the biological validity of VIP-OT–inferred pairings, we applied the method to a previously published single-cell FTIR dataset^25^ comprising MDA-MB-231 human breast adenocarcinoma cells treated with 16 drugs spanning 9 mechanisms of action (MoAs) at IC_50_ concentrations. A complete list of drugs, MoAs, and IC_50_ values is provided in **Supplementary Table 1**. Conceptually, these MoAs broadly fall into two groups: general metabolic inhibitors and targeted pathway/protein inhibitors. Against this diverse perturbation landscape, each VIP-OT perturbation vector represents the spectral displacement between a treated cell and its assigned control counterpart in the high-dimensional vibrational space.

Uniform Manifold Approximation and Projection (UMAP)^44^ of per-cell perturbation vectors showed clear separation between different drug MoAs, with treatments forming distinct clusters (**Fig. 2a**). Consistently, hierarchical clustering analysis (HCA) of mean perturbation vectors clearly grouped drugs sharing the same MoA, annotated with similar color hues (**Fig. 2b**). These results indicate that VIP-OT pairings preserve drug-specific response patterns at both single-cell and condition levels.

**Fig. 2.**
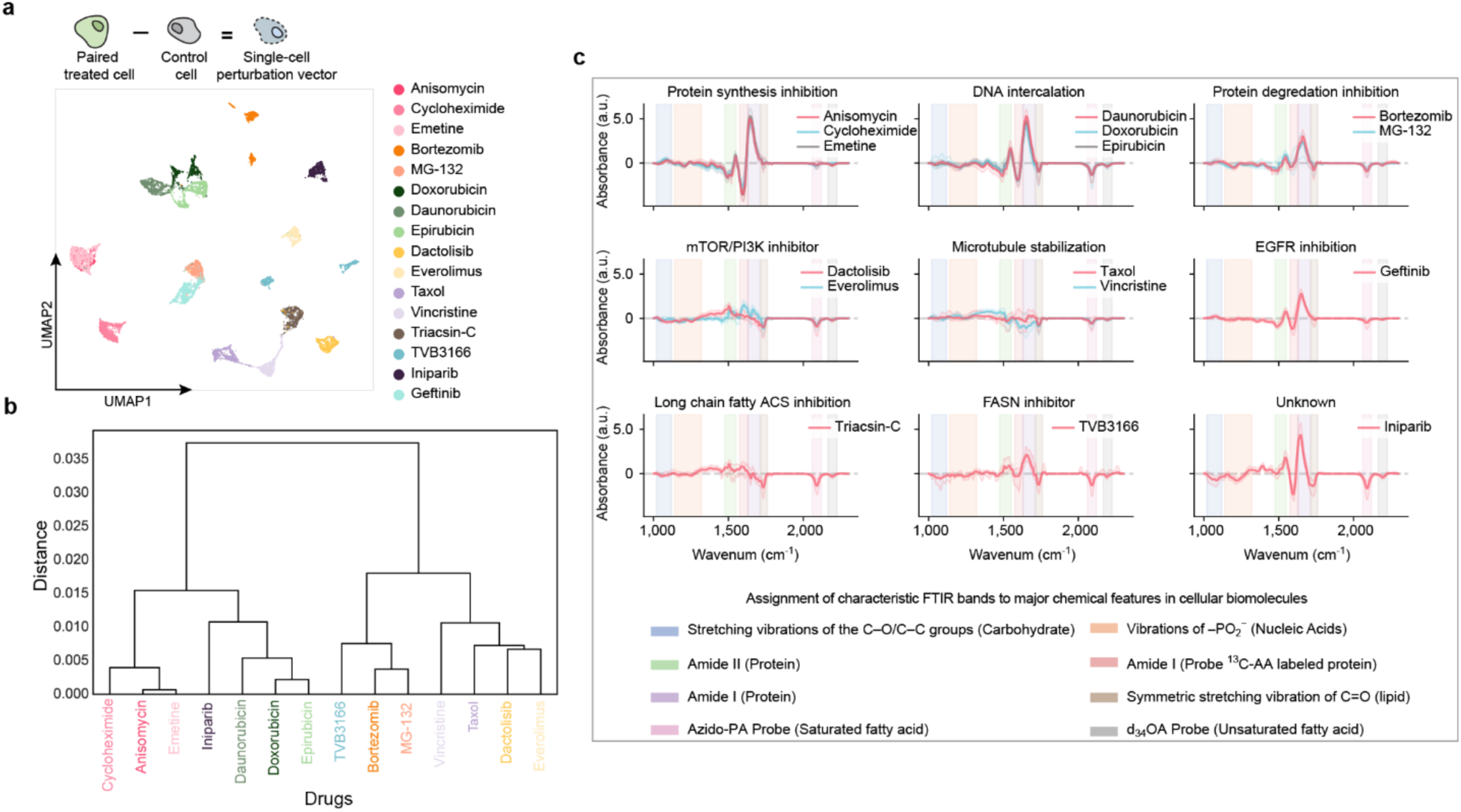
Biological relevance of VIP-OT–inferred perturbation vectors. **a,** UMAP projection of single-cell perturbation vectors derived from VIP-OT. Each point represents a single perturbed cell, and colors indicate drug treatment. **b,** Hierarchical clustering of average perturbation vectors shows drugs grouping by known MoAs. **c,** Difference Feature Spectra (DFSP) for representative MoAs. Traces show mean spectral shifts between paired cells; shaded regions indicate 95% confidence intervals. Colored regions mark characteristic vibrational modes of biomolecules and metabolic probes.

To assess biochemical interpretability, we computed *Difference Feature Spectra* (DFSP) by subtracting each treated spectrum from its paired control spectrum. Because VIP-OT establishes cell-to-cell correspondences, these difference spectra directly capture treatment-induced shifts at single-cell resolution, where upward and downward peaks indicate relative increases or decreases in absorbance. For example, anisomycin, cycloheximide, emetine-treated cells—subject to protein synthesis inhibition^45–47^—showed similar spectral patterns and marked reductions in the ^13^C-amino acid–labeled Amide I region, consistent with suppressed incorporation of labeled amino acids into newly synthesized proteins (**Fig. 2c**). Iniparib, an experimental anticancer drug once claimed to be a poly (ADP-ribose) polymerase (PARP) inhibitor^48^ but whose specific mechanism remains unknown^49^, clustered closer to DNA-targeting drugs such as daunorubicin, doxorubicin, and epirubicin (**Fig. 2c**).

Additional analyses further support the plausibility of OT-based correspondences (**Supplementary Figs. S1–S4**). Feature-level UMAPs (**Supplementary Fig. S1**) revealed MoA-consistent suppression or enhancement of probe- and biomolecule-associated spectral bands. Heatmaps of averaged difference spectra across probe-sensitive and endogenous regions (**Supplementary Fig. S2**) captured distinct biochemical fingerprints for each drug. Metabolic coordinate plots (**Supplementary Fig. S3**) and corresponding per-axis boxplots (**Supplementary Fig. S4**) mapped drug-specific alterations in protein synthesis, saturated fatty acid metabolism, and unsaturated fatty acid metabolism, highlighting both shared and divergent biosynthetic effects within MoA classes.

Together, these results demonstrate that VIP-OT–inferred perturbation vectors faithfully encode biologically meaningful, MoA-specific metabolic shifts, establishing a robust foundation for downstream analyses.

### VIP-OT enables retrospective, single-cell tracing of baseline-dependent response heterogeneity

Drug treatments often elicit heterogeneous responses across individual cells within a population^50^, and such variability has been recognized as a major factor influencing therapeutic efficacy and resistance^51^. A central hypothesis in pharmacology is that a cell’s baseline metabolic state predetermines its sensitivity to a given drug^52–54^, yet testing this at the single-cell level has been hindered by the lack of paired pre- and post-treatment data. To investigate the hypothesis, we utilized the retrospective alignment capability of VIP-OT. This framework enables a direct, single-cell analysis of how pre-existing molecular composition shapes therapeutic response.

As a case study, we applied this approach to dissect the heterogeneous response of MDA-MB-231 human breast adenocarcinoma cells to anisomycin, an antibiotic that inhibits protein synthesis by binding to ribosomes and activating stress-responsive MAPK pathways^55^. Anisomycin has also been reported to influence lipid metabolism through indirect regulation of lipid-associated signaling pathways and energy homeostasis^56^. Given these mechanisms, we reasoned that anisomycin’s effects might vary according to a cell’s baseline lipid content. We quantified three metabolic activities (Methods) and found that saturated fatty acid (SFA) uptake distribution displayed a marked broadening under treatment. VIP-OT was used to align control and anisomycin snapshots.

To validate the OT mapping, we compared each cell’s baseline value to both the OT-estimated barycentric outcome and the displacement. Baseline values strongly tracked expected post-treatment value (Spearman ρ = 0.879; slope = 1.03) but not the displacement (Spearman ρ = 0.068; slope = 0.029), indicating rank preservation without baseline-dependent shifts (**Supplementary Fig. S5**). We then visualized the OT-weighted joint distribution of SFA uptake (**Fig. 3a, left**). Relative to the identity line, the density exhibited a coherent displacement and pronounced broadening, consistent with structured response heterogeneity. To expose this structure, we applied k-means clustering^57^ (k = 2) to the treated metric and overlaid the subgroup labels on the joint distribution (**Fig. 3a, right**). This analysis identified two well-separated strata: high SFA metabolism group and low SFA metabolism group of comparable size (Low-SFA group, *n* = 116; High-SFA group, *n* = 129; silhouette score^58^ = 0.612) (**Fig. 3b**).

**Fig. 3.**
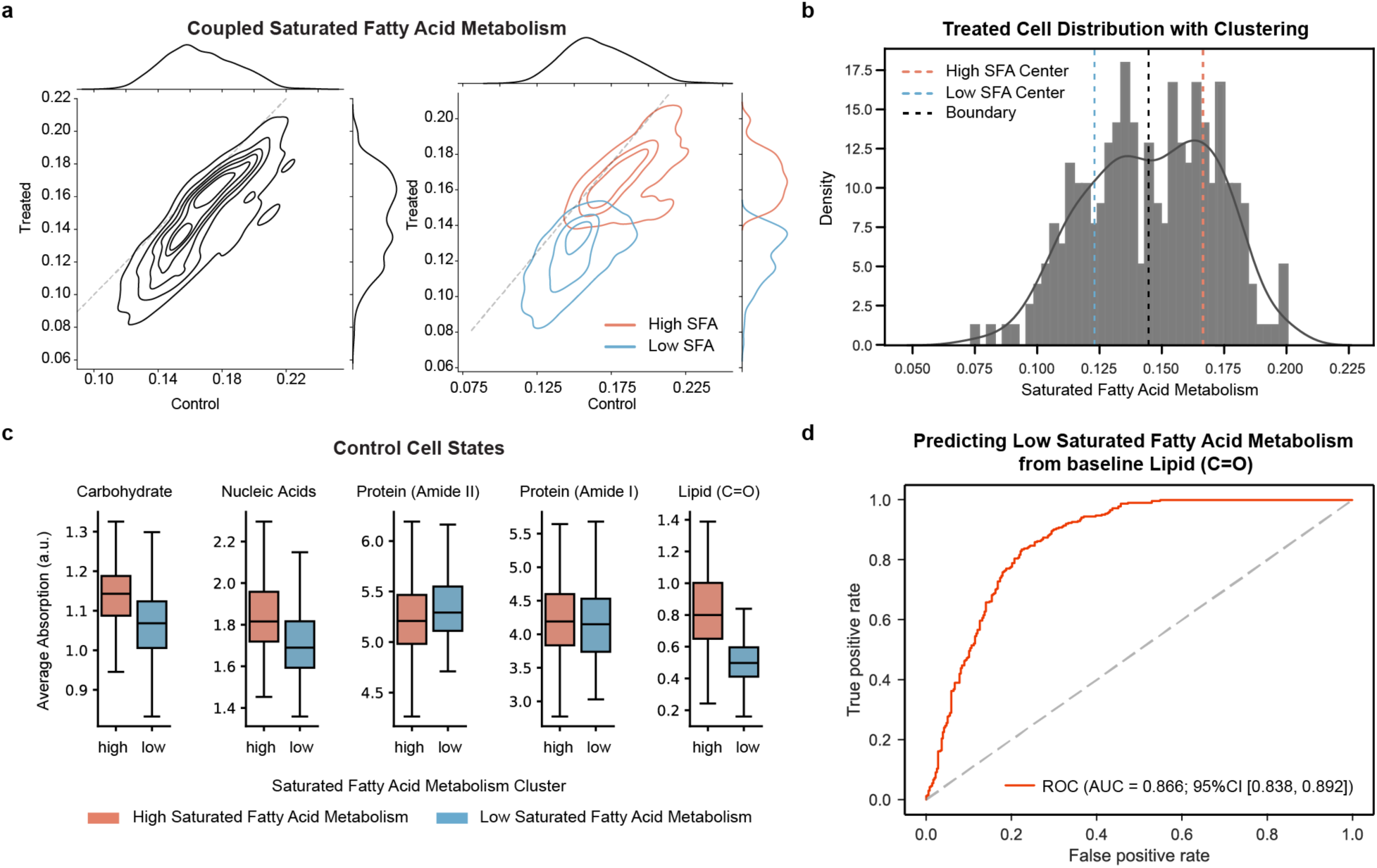
VIP-OT enables retrospective tracing of baseline-dependent response heterogeneity. **a, Left** OT-weighted joint kernel density of saturated fatty acid (SFA) metabolism between anisomycin-treated cells and control cells. **Right:** k-means clustering (k = 2) stratified treated cells into high-SFA and low-SFA subgroups (Low-SFA group, *n* = 116; High-SFA group, *n* = 129; silhouette score= 0.612); subgroup labels were propagated to control counterparts via VIP-OT pairing. **b,** Distribution of treated cells with clustering centers and the midpoint boundary (centers: Low-SFA = 0.123, High-SFA= 0.167; midpoint = 0.145). **c,** Baseline absorbance across selected vibrational bands in control cells stratified by retrospective subgroup assignment. Cells later classified as Low-SFA exhibited reduced lipid-associated absorbance (C=O stretch) than those classified as High-SFA (Mann–Whitney test, U = 15,715; p = 1.47 × 10^−61^; Cliff’s δ = −0.73). **d,** ROC analysis using baseline ester to predict Low-SFA assignment (AUROC = 0.866; 95% CI [0.838, 0.892]). Labels on control assigned by weight-proportional sampling from OT couplings.

We next propagated these subgroup labels back to control cells using OT-derived couplings and examined their baseline vibrational spectra. Notably, control retrospectively assigned to the low-SFA subgroup showed reduced absorbance in the ester band (∼1740 cm⁻¹) relative to the high-SFA subgroup (Mann– Whitney test, p = 1.47 × 10^−61^; Cliff’s δ = −0.73) (**Fig. 3c**). This result supports a model in which cells with lower baseline lipid ester content are more vulnerable to anisomycin-induced suppression of SFA uptake, illustrating the capacity of VIP-OT to uncover causal links between pre-treatment biochemistry and drug sensitivity.

Finally, we evaluated the predictive value of baseline lipid composition directly. A prospective test using the ester band signal alone achieved strong discrimination of future subgroup assignment (AUROC = 0.866; 95% CI, 0.838–0.892) (**Fig. 3d**). Together, these analyses demonstrate that single-cell vibrational baselines carry predictive information about anisomycin response heterogeneity, validating VIP-OT as a powerful framework for dissecting the metabolic determinants of heterogeneous drug efficacy.

### VIP-OT enables prediction of single-cell drug responses from pre-treatment vibrational spectra

Intra-tumor heterogeneity (ITH) poses a major challenge to effective therapy, as different subpopulations within the same tumor often respond divergently to treatment^59^. A central goal in personalized medicine is to predict heterogeneous drug responses at the single-cell level directly from pre-treatment cellular states^60,61^. Building on the pseudo-pairings established by VIP-OT, we developed a computational framework to achieve this, enabling the prospective prediction of post-treatment vibrational spectra from unperturbed baseline measurements.

The framework operates on a ‘guilt-by-association’ principle: cells with similar baseline biochemical features tend to exhibit similar perturbation signatures^62–66^. For each drug condition, OT couplings yield barycentric treated targets for every control cell. We then trained drug-specific models *f*^(*d*)^: *x* → *ŷ*, that map baseline spectra *x* to their predicted treated states *ŷ* for each drug *d*, individually. To preserve heterogeneity, we adopted a nearest-neighbor–based strategy: each test cell is matched to its most similar training control cell in the spectral space, and the associated OT-weighted treated average is transferred as its predicted response (Methods). This design leverages a shared manifold of control cells across drugs, while allowing predictions to remain cell-specific and drug-dependent.

We benchmarked the framework on a panel of 10 drugs spanning diverse MoAs^25^, all sharing the same pool of 1,084 baseline control cells. This unified setting allowed for consistent cross-drug evaluation. For each drug, the model predicts the full post-treatment vibrational spectrum for an unperturbed cell from its baseline spectrum, and derives three functional metabolic readouts (protein-synthesis rate, saturated fatty-acid metabolism and unsaturated fatty-acid metabolism) at single-cell resolution. Across 20-fold cross-validation, predicted spectra closely matched experimental measurements, with median Pearson correlations exceeding 0.99 for all drugs (**Fig. 4a**). Functional readouts were recovered with similarly high fidelity: protein synthesis rates were predicted with high accuracy (R² ranging from 0.805 to 0.930) and narrow MAE/MSE distributions across drugs (**Fig. 4b**). Saturated and unsaturated fatty acid uptake were similarly well captured (**Supplementary Fig. S6**). To enable direct cross-drug comparison, we defined a composite prediction performance score combining average spectral R², spectral correlation, and metabolic R² (Methods). This revealed drug-specific predictability ranging from 0.945 (cycloheximide) to 0.847 (everolimus) (**Fig. 4c**), consistent with the biochemical complexity of their mechanisms.

**Fig. 4.**
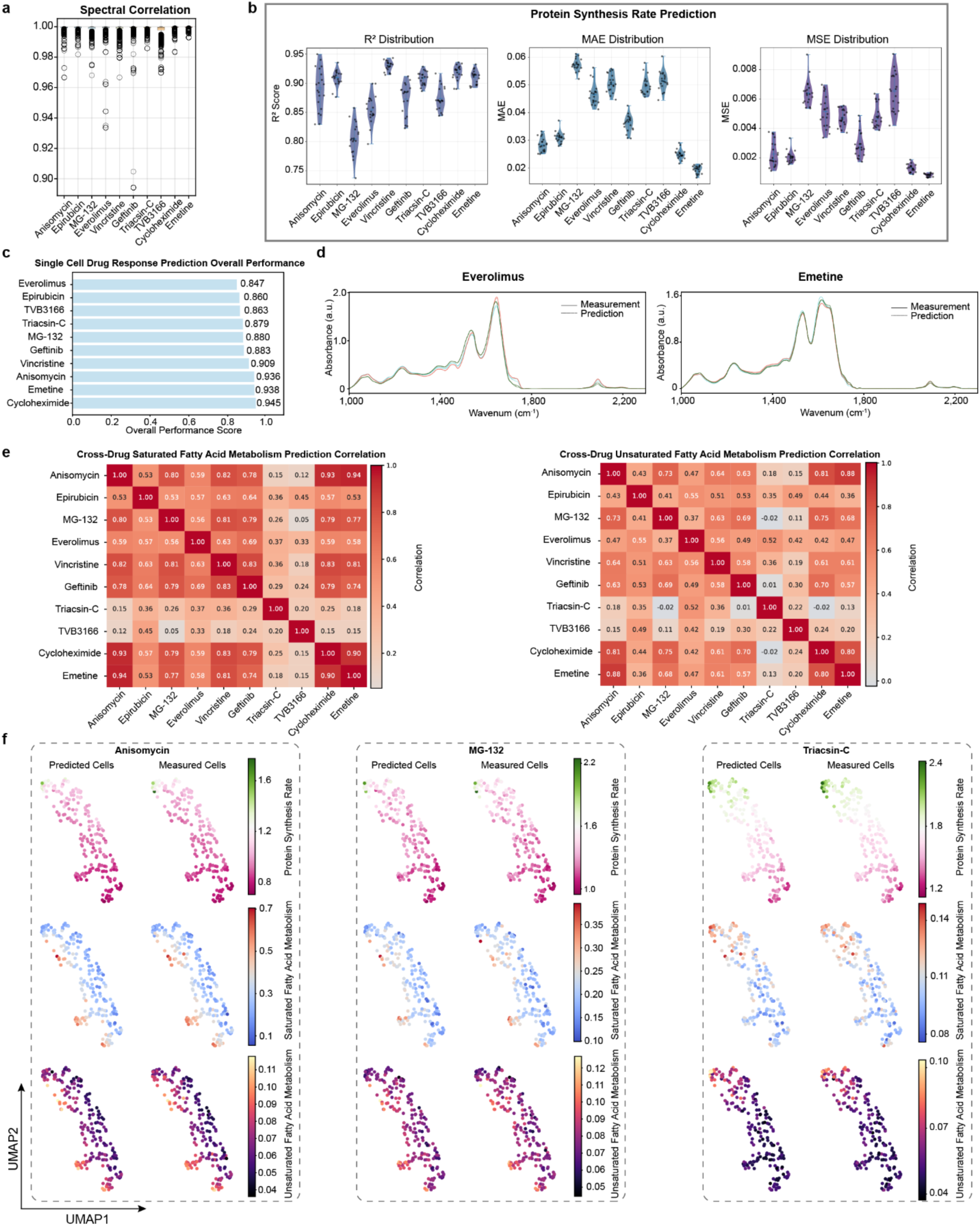
VIP-OT enables drug-specific prediction of single-cell metabolic responses. **a,** Distributions of Pearson correlation between predicted and measured vibrational spectra across 20 cross-validation folds for each drug; high correlations indicate accurate spectrum-level predictions. **b,** Violin plots showing R², MAE, and MSE distributions of protein synthesis rate predictions across drugs and validation folds; other metabolic readouts (saturated and unsaturated fatty acid uptake) are shown in Supplementary Fig. S6. **c,** Overall prediction performance score per drug, defined as the mean of spectral R², spectral correlation, and average metabolic R² across three activities. Drugs differ in predictability, with scores ranging from 0.847 to 0.945. **d,** Example spectral overlays for three representative test cells treated with different drugs. Solid lines: experimental spectra; dashed lines: predicted spectra. **e,** Cross-drug correlation heatmaps of predicted fatty acid metabolic responses. Drugs with similar MoAs show higher correlation. **f,** UMAP projection of test control cells colored by predicted and measured values of three metabolic activities under three drugs. Prediction patterns match measured patterns at single-cell resolution.

Predicted spectra were nearly indistinguishable from experimental measurements. For representative test cells across two drugs, the overlaid spectra (predicted: dashed; measured: solid) showed almost perfect coincidence across the entire vibrational fingerprint (**Fig. 4d**). This highlights the ability of the model to recover full spectral information, not merely coarse features or summary metrics. Furthermore, the predicted metabolic readouts also captured drug-specific phenotypic features: correlation matrices of fatty acid metabolism responses revealed known MoA relationships (**Fig. 4e**). For example, anisomycin and emetine (both protein synthesis inhibitors) displayed the highest correlation in predicted responses

(correlation = 0.84), whereas MG-132 (proteasome inhibitor) and TVB-3166 (FASN inhibitor) exhibited minimal similarity (correlation = 0.12) (**Fig. 4e**). These results show that the predictive framework captures mechanistic convergence and divergence directly from baseline vibrational states, in full agreement with biological expectation. Extending to the single-cell population level, predicted phenotypic landscapes mirrored the true metabolic readouts with striking fidelity. UMAP projections of predicted metabolic activities under three representative drugs reproduced the fine-grained heterogeneity within each population, closely matching the experimental embeddings (**Fig. 4f**).

Together, these results demonstrate that VIP-OT transforms vibrational imaging into a predictive framework, enabling accurate, drug-specific modeling of single-cell responses directly from baseline metabolic states. This predictive capacity underscores that baseline vibrational spectra contain rich biochemical information which, when coupled with VIP-OT-derived mappings, can anticipate treatment outcomes at single-cell resolution. Extending this concept, virtual perturbation profiling of patient-derived baseline spectra could enable in silico “treatment” at the single-cell level, offering a potential path toward personalized therapy in heterogeneous tumors.

### Dynamic trajectory modeling with spectral velocity (SVL) uncovers dose-dependent responses

We next extended the VIP-OT framework to model dynamic cellular processes by reconstructing dose-dependent response trajectories from a series of static snapshots. To achieve this, we introduce *Spectral Velocity* (SVL), a vector descriptor that quantifies the magnitude and direction of a cell’s metabolic state transition between adjacent doses. By applying OT to couple cell populations across a graded drug concentration series, we can compute these vectors for each cell, assembling a dynamic map of the population’s response trajectory.

To demonstrate this capability, we profiled MDA-MB-231 human breast adenocarcinoma cells treated with anisomycin (protein synthesis inhibitor) and TVB-3166 (FASN inhibitor) across graded concentrations, from subthreshold to cytotoxic concentrations (Methods; **Supplementary Table 2**). These two drugs are chosen as representative inhibitors of protein synthesis and fatty-acid metabolism, respectively, thereby probing distinct biochemical axes of diverse perturbations. At the population level, OT-derived couplings revealed coherent, dose-dependent spectral remodeling (**Fig. 5a**). For anisomycin, low doses produced sharp suppression of the ^13^C-labeled amide I band and reciprocal increases in native amide I, consistent with protein synthesis inhibition, whereas high doses caused broad attenuation across nucleic acids, carbohydrates, and lipids signals, reflecting a collapse of global metabolism. In contrast, TVB-3166 induced coherent lipid-specific suppression at intermediate doses but gave rise to diffuse and noisy changes at high concentrations, likely reflecting disorganized cellular states under excessive cytotoxicity. The metabolic variations of treated cells could influence the activation of key signaling pathways, ultimately affecting downstream biological processes. Beyond average spectral shifts, VIP-OT also reveals how biochemical heterogeneity evolves across doses. Kernel density estimates of absorbance distributions in key biomolecular regions (**Fig. 5b**) showed that anisomycin-treated cells maintained relatively stable carbohydrate distributions at low dosage, but exhibited broadening and down-shifting at high dosage. These spectral shifts are consistent with stress-induced metabolic divergence, in line with the Integrated Stress Response (ISR) pathway, which suppresses global protein synthesis and alters intracellular carbohydrate composition^67^. TVB-3166, by comparison, showed saturated fatty acid uptake converged under high dosage, indicating uniformly suppressed lipid metabolism due to FASN inhibition and reduced demand for exogenous fatty acids^68^. Moreover, cells treated by TVB-3166 produced peak heterogeneity at 10 μM followed by collapse at 200 μM, reflecting variable mid-dose responses and uniform shutdown at extreme stress.

**Fig. 5.**
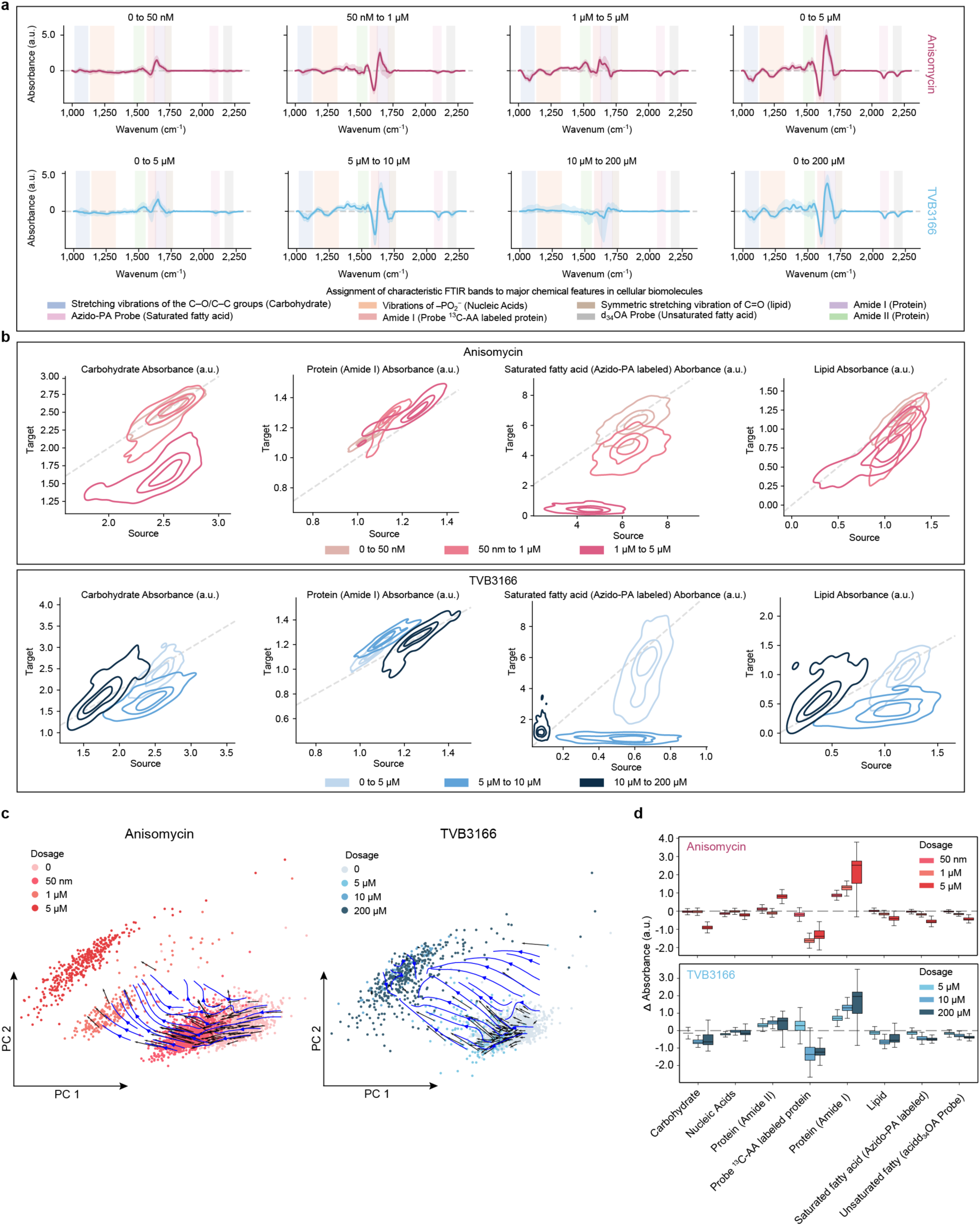
Spectral Velocity Maps reveal dose-resolved biochemical trajectories and evolving single cell heterogeneity. **a**, DFSP between adjacent dose levels reveal progressive molecular perturbations along the dosage continuum. **b**, Kernel density plots of OT-coupled single-cell absorbance values across selected spectral bands highlight dose-dependent redistribution of biochemical heterogeneity. **c**, Spectral Velocity Maps constructed from OT-derived displacement vectors visualize directional transitions between dose levels in a shared low-dimensional embedding (e.g., PCA). Black arrows represent sampled displacement vectors from high-confidence OT pairings (≥ 0.3), and blue streamlines represent interpolated velocity fields. These trajectories indicate the magnitude and direction of spectral shifts at single-cell resolution. **d**, Boxplots summarize absorbance differences across doses within characteristic spectral regions.

Moving beyond population-level trends towards interpretable single-cell dynamics, we for the first time introduce *Spectral Velocity* (SVL)—a novel descriptor quantifying the direction and magnitude of each cell’s transition between adjacent doses. Conceptually, spectral velocity estimates where a cell is “headed” in spectral state space, and describes the minimal transitions required for each cell to reach its most probable next state, effectively predicting its future state. Computed in a shared low-dimensional embedding (here PCA, though the approach is embedding-agnostic), velocity vectors capture cell-specific rates and directions of change (Methods). Spectral velocity maps revealed striking drug-dependent differences (**Fig. 5c**). Under anisomycin, vectors formed a smooth, directed flow from low to high doses, indicating a coherent biochemical trajectory with varying speeds across cells. In contrast, TVB-3166 produced a fragmented velocity field, with disorganized or abruptly terminating vectors at cytotoxic doses, reflecting disrupted metabolic progression. These vector fields provide a high-resolution visualization of how heterogeneous cell populations traverse metabolic state space in response to perturbation.

We further validated this trajectory framework by performing spontaneous Raman spectroscopy measurements on the same anisomycin and TVB-3166 dosage series (Methods; **Supplementary Table 3**). Raman spectroscopy, as a complementary vibrational modality with higher spectral resolution than most FTIR spectra, reproduced the FTIR-inferred dose-dependent patterns and VIP-OT–derived trajectories (**Supplementary Fig. S7–S8**). This cross-platform consistency demonstrate that VIP-OT generalizes across vibrational platforms, extending its applicability to Raman spectroscopy and broadening its relevance for multimodal biochemical phenotyping.

Together, these analyses advance vibrational imaging from static description to dynamic trajectory modeling. The introduction of Spectral Velocity provides a quantitative and generalizable framework for tracing how cellular states evolve along a continuum, bridging static snapshots with predictive models of dynamic biological processes.

### Path-resolved decomposition reveals hidden asymmetry in drug combination effects

Combination therapies are a cornerstone of cancer treatment, aiming to improve efficacy and prevent resistance^69^. Conventional analysis of combination therapies classifies drug interactions based on final cellular endpoints, such as viability^70–72^. However, these endpoint scores neglect the molecular routes, obscuring whether different treatment paths—for instance, adding drug A then B, versus B then A— converge on the same phenotype through distinct mechanisms. In reality, many clinically relevant phenomena are known to be path-dependent, where distinct perturbation routes can converge on a shared state through different biochemical transition^73,74^. We therefore adapted the VIP-OT framework to move beyond endpoint analysis and perform a path-resolved decomposition of drug combination effects at the single-cell level.

To test this framework, we treated MDA-MB-231 human breast adenocarcinoma cells with Bortezomib (proteasome inhibitor) and Gefitinib (EGFR inhibitor), two drugs with distinct MoAs, across nine conditions encompassing IC_20_ and IC_50_ single-agent as well as their matched combinations (Methods; **Supplementary Table 4**). For each combination, we computed OT couplings from each preceding single-agent condition to the final combination state (**Fig. 6a**). This approach allowed us to reconstruct and compare the distinct molecular paths leading to the same outcome.

**Fig. 6.**
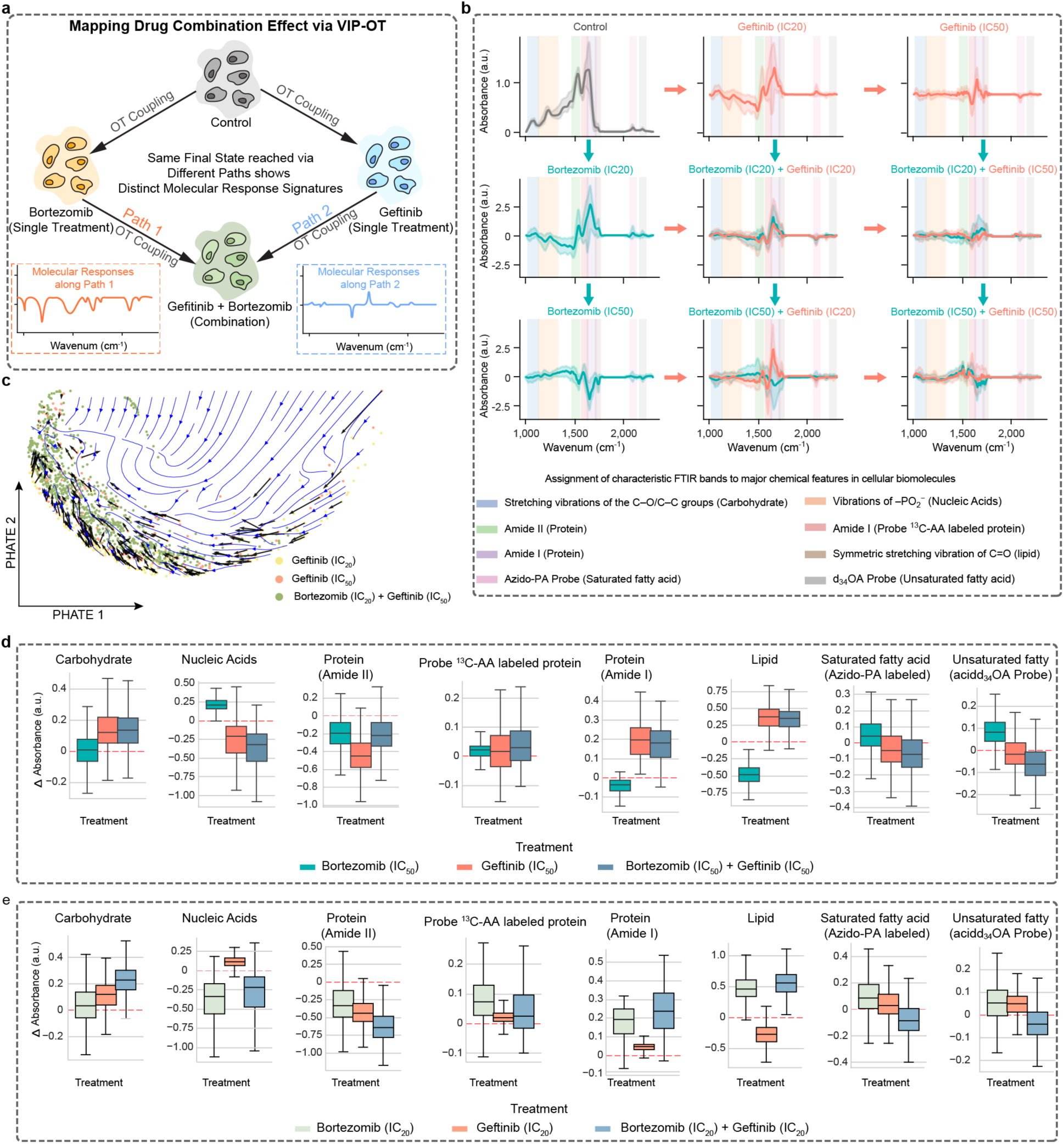
Path-resolved decomposition of drug combination responses via VIP-OT. **a**, Schematic of path-aware OT analysis, where spectral transitions are independently mapped from each single-agent condition to the combination state, enabling reconstruction of distinct molecular trajectories. **b,** DFSP derived from OT pairings reveal asymmetric biochemical transitions into the combination state. Each path contributes distinct spectral changes, indicating that similar endpoints can arise via divergent molecular mechanisms. Blue denotes increasing bortezomib at the current gefitinib level; orange denotes increasing gefitinib at the current bortezomib level. The top-left panel shows raw control spectra. **c,** Spectral Velocity Map constructed in PHATE space^78^. Black arrows represent OT-derived displacement vectors sampled from high-weight cell pairs; blue streamlines denote interpolated velocity fields, capturing coherent pseudo-temporal flow along treatment progression. **d–e**, Boxplots summarize average spectral differences across key molecular regions under high-dose (d) and low-dose (e) treatment conditions. Combination treatments exhibit the strongest suppression in lipid-associated vibrational band.

The path-resolved analysis revealed a marked asymmetry in the underlying molecular transitions (**Fig. 6b**), in each DFSP, blue path denotes adding bortezomib at the current gefitinib level, and orange path denotes adding gefitinib at the current bortezomib level. For example, in the Bor IC_50_ + Gef IC_20_ group, the orange path (Gefitinib added on top of Bor IC_50_) showed a pronounced decrease in ^13^C-amide I together with an increase in native amide I, indicating suppressed protein synthesis rate. This aligns with EGFR blockade acting through the PI3K–AKT–mTORC1 axis, where inhibition of 4EBP1 and S6K phosphorylation disrupts cap-dependent translation initiation^75,76^. In contrast, the blue path (raising Bortezomib from IC_20_ to IC_50_ on a Gef IC_20_ background) displayed a more global decrease in protein-associated signals, consistent with proteasome inhibition driving proteotoxic stress and compensatory autophagy to maintain proteostasis^67,77^.

Although the cells reached a similar endpoint, the molecular paths taken were demonstrably different, highlighting a mechanistic asymmetry invisible to conventional synergy analysis. This dynamic progression was further visualized by embedding cells in PHATE space^78^, where spectral velocities showed coherent trajectories flowing smoothly into the final combination state (**Fig. 6c**).

Quantitative summaries across vibrational bands showed that combination treatments consistently produced the strongest suppression of lipid-associated signals, beyond either monotherapy alone (**Fig. 6d–e**). This compounded suppression indicates that co-treatment perturbs lipid metabolism more strongly than either agent alone, likely reflecting synergistic interference between proteostasis disruption and EGFR signaling blockade. Notably, this pattern aligns with prior reports that proteasome inhibition enhances apoptosis in EGFR-targeted contexts, supporting a convergent pressure on lipid biosynthesis pathways^79^.

These results establish VIP-OT as a tool for path-aware analysis of drug combinations. By disentangling how different treatment routes contribute to a shared endpoint, the framework shifts synergy studies from static summaries to dynamic, path-resolved decompositions. This capability provides a new front for designing more precise and rational combination therapies by considering not just the destination but the journey a cell takes to get there.

## Discussion

Achieving a predictive understanding of cellular behavior requires methods that can model heterogeneous and dynamic responses to perturbations. A major challenge for many powerful single-cell profiling techniques, including vibrational imaging, is that they yield static population snapshots, fundamentally limiting their ability to capture the transitions between cellular states. In this study, we address this gap by introducing VIP-OT, a framework that integrates multiplexed vibrational imaging with optimal transport to computationally reconstruct these missing cellular dynamics. The central advance of VIP-OT is its ability to transform vibrational imaging from a descriptive to a predictive and dynamic modality. By establishing soft correspondences between untreated and perturbed cells in spectral space, VIP-OT transforms static single-cell measurements into dynamic, predictive models of response. This platform is capable of classifying MoAs, tracing baseline heterogeneity, predicting drug responses, reconstructing dynamic trajectories, and resolving path-dependent effects at the single-cell level.

Through inferred perturbation vectors, VIP-OT classifies drugs with similar of distinct MoAs. Iniparib, whose specific mechanism remains unknown, clustered closer to DNA-targeting drugs, indicating a potential MoA. By retrospectively pairing treated cells with their inferred control counterparts, VIP-OT makes it possible to trace treatment outcomes back to pre-existing biochemical states. This capability revealed that baseline lipid-associated ester band intensity stratifies anisomycin responses: cells with lower lipid content were more strongly suppressed in saturated fatty acid metabolism. VIP-OT, for the first time, enables per-cell attribution of heterogeneous responses to baseline metabolic features, which is a task previously inaccessible in vibrational imaging. Meanwhile, we acknowledge that the baseline cellular heterogeneity and states represent a continuum, we chose K=2 for the purpose of better and easier biologic interpretation. Building on this correspondence, we developed a non-parametric prediction framework that accurately forecasts single-cell drug responses from unperturbed spectra, achieving near-perfect correlations across both full spectra and functional metabolic readouts. A further innovation is the introduction of spectral velocity, a vectorial descriptor of per-cell state transitions during dynamic process. Velocity fields captured coherent, directed trajectories under anisomycin (protein synthesis inhibitor) but fragmented dynamics under TVB-3166 (FASN inhibitor), reflecting the difference between ordered stress responses and disorganized collapse. This moves vibrational profiling beyond endpoint classification toward dynamic modeling of dose-dependent and temporal responses. Finally, VIP-OT enables decomposition of combination treatments into path-specific molecular routes. By separately coupling each single-agent state to the combination outcome, we revealed asymmetric trajectories: bortezomib (proteasome inhibitor) and gefitinib (EGFR inhibitor) converged on a shared endpoint but through distinct biochemical transitions, consistent with their known MoAs. Such path-resolved views expose asymmetries invisible to conventional bulk synergy scores.

VIP-OT can further evolve in several notable directions. First, our present work is based on immortalized cell line models. A critical next step is expanding VIP-OT to 3D organoids and patient-derived *in vitro* models, where microenvironmental context and spatial heterogeneity are preserved. Achieving this requires higher-throughput vibrational imaging: discrete frequency infrared (DFIR) imaging offers 10-100× faster acquisition than FTIR^80–82^, and stimulated Raman scattering can drastically accelerate chemical imaging over the conventional Raman microscopy^19,83^. Second, although we focused on chemical perturbations, the VIP-OT framework is fundamentally agnostic to perturbation type and readily generalizes to other perturbation types. It can be applied to CRISPR-based genetic perturbations or transcription factor modulation^84–86^, providing a way to chart convergent and divergent routes of cellular reprogramming beyond drug screening. Third, analytical principles of VIP-OT are modality-agnostic and naturally extend to multimodal integration. Vibrational phenotypes can be co-analyzed with single-cell or spatial transcriptomics, linking metabolic states to gene-regulatory programs^87^. We recently showed that vibrational spectra can be co-analyzed with transcriptomic data^88^. VIP-OT framework provides a natural bridge for such systematic integration. Finally, VIP-OT points toward a translational vision: baseline vibrational profiles from a patient biopsy could be used to simulate responses to multiple drugs or combinations at single-cell resolution. This would enable early stratification of therapeutic options, identification of resistant subpopulations, and rational design of personalized regimens. In this sense, VIP-OT represents a concrete step toward the AI Virtual Cell, where predictive virtual perturbations guide precision medicine.

## Methods

### Cell Line and Materials

MDA-MB-231 cells (ATCC HTB-26) was purchased from ATCC. For reagents, azido-palmitic acid (1346) was purchased from Click chemistry tools; algal amino acid mixture (U-13C, 97-99%, CLM-1548) was purchased from Cambridge; deuterated oleic acid (683582) was purchased from Sigma-Aldrich. For drugs, anisomycin (A9789), bortezomib (179324-69-7), gefitinib (184475-35-2) and TVB-3166 (SML1694) were purchased from Sigma-Aldrich. For cell culture agents, DMEM medium (11965), FBS (10082), penicillin/streptomycin (1514), were purchased from ThermoFisher Scientific. CaF_2_ substrates (CAFP13-1) were purchased from Crystran.

### Probe preparation and media recipe for cell labeling

#### Azido palmitic acid-bovine serum albumin (BSA) solution

For the solution, couple azido-palmitic acid with BSA to prepare a 2 mM stock solution. Prepare 20 mM sodium palmitic acid solution by dissolving palmitic acid in NaOH solution with the following recipe: azido-PA (5.5 mg) + 1.0 ml dd-H_2_O + 35 μl 1 M NaOH. Mix and incubate the solution in 70 °C water baths until no oil droplets are visible. Then slowly add the sodium palmitic acid solution into 2.7 ml 20% BSA under room temperature water baths. Quickly add 6.3 ml DMEM culture medium and filter the solution with a 0.22 μm sterile filter.

#### 13C-amino acids DMEM

Here, 4 mg ml^-1^ algae ^13^C-amino acids mix was dissolved in dd-H_2_O with 10% FBS and 1% penicillin, which matched the concentrations of regular amino acids in DMEM.

#### Deuterated oleic acid-bovine serum albumin (BSA) solution

For the solution, couple d_34_-oleic acid with BSA to prepare a 2 mM stock solution. Prepare 20 mM oleic acid solution by dissolving oleic acid in NaOH solution with the following recipe: d_34_ oleic acid (6.3 mg) + 1.0 ml dd-H_2_O + 24 μl 1 M NaOH. Mix and incubate the solution in 70 °C water baths until no oil droplets are visible. Then slowly add the d_34_ oleic acid solution into 2.7 ml 20% BSA under room temperature water baths. Quickly add 6.3 ml DMEM culture medium and filter the solution with a 0.22 μm sterile filter.

#### Cell culture

MDA-MB-231 cells were cultured in DMEM media supplemented with 10% FBS and 1% penicillin. Cells were grown in a humidified atmosphere containing 5% CO_2_ at 37 °C in the incubator. At ∼80% confluence, cells were dissociated with trypsin and passaged.

#### Sample preparation for drug-treated cells with labeling

MDA-MB-231 cells were seeded on clean CaF_2_ substrates with 50,0000 cells per well in cell culture media (DMEM, 10% FBS, 1% penicillin) overnight for control and other drug treatment conditions. Then the culture media was replaced by ^13^C-amino acids DMEM with 50 M azido-palmitic acid, 50 M d_34_ oleic acid, either single drug at chosen concentrations. Drugs were prepared in 100% DMSO and diluted to 0.1% DMSO in labeling media. For the control group, only cell labeling media with 0.1% DMSO was added (without any drugs). Cells were treated for 48 hrs. The culturing time was selected based on the trade-off between signal and experimental time. After that, cells were fixed by 4% PFA at room temperature for 15 min and washed three times with PBS buffer and five times with dd-H_2_O. The samples were then air-dried before FTIR imaging.

#### FTIR imaging

Agilent Cary 620 Imaging FTIR equipped with an Agilent 670-IR spectrometer and 128 × 128-pixels FPA mercury cadmium telluride (MCT) detector was used in the transmission mode. A background spectrum was collected on a clean CaF_2_ substrate using 256 scans at 8 cm⁻¹ spectral resolution, suggesting that the IR absorbance was measured every 4 cm⁻¹. Cell spectra were recorded using 128 scans at 8 cm⁻¹ spectral resolution. A ×25 IR objective (pixel size, 3.3 μm, 0.81 numerical aperture (NA)) was used for cell imaging.

#### Spontaneous Raman imaging

Spontaneous Raman imaging was performed using an upright confocal Raman microscope (Xplora, HORIBA Jobin Yvon). Cell samples were illuminated by 532 nm laser (80 mW on sample) through a ×50 objective (air, NA 0.75, MPlan N, Olympus). Raman images were acquired using the point-by-point mapping mode with an acquisition time of 5s and 1× accumulation for each point measurement. The step size was set as 7 μm. The grating was set as 1200 gr/mm. Both the slit size and the hole size was set as 100 μm.

#### FTIR data preprocessing

FTIR data preprocessing was performed using both the commercial software Cytospec and home-built MATLAB(MathWorks) scripts with the following steps: (1) PCA noise reduction to denoise and reconstruct the spectra, the top 30 principal components were kept for imaging data collected from each condition, including control and drug treatment; (2) quality test to remove pixels with a low signal-to-noise ratio in both the fingerprint region and the cell-silent region; (3) rubber-band baseline correction for spectral correction; (4) single-cell segmentation on cell images using CellProfiler^89^; (5) single-cell spectrum extraction applying generated single-cell masks on the processed FTIR imaging data and (6) signal normalization (normalizes the area of the whole spectra to 1) was applied on each single cell spectrum.

#### Raman data processing

Raw Raman spectra were first processed in the LabSpec 6 software (HORIBA). The “Despike” function was used to remove cosmic rays. The “Threshold” function was used to remove spectra with high background. The “Correction” function was used to subtract background from raw spectra by selecting a field of view without any cells. After that the spectral range from 2830 cm^-1^ to 3000 cm^-1^ was integrated to generate a cell image, using custom-written MATLAB scripts. The cell image was used to segment single cells and generate a cell segmentation mask with the Otsu thresholding method using the CellProfiler software. Based on the cell segmentation mask, single-cell spectra were generated by averaging all the spectra within each cell, using custom-written MATLAB scripts. CH signal normalization (normalizes the area of the (a)symmetric CH stretching (2815 cm⁻¹-3015 cm⁻¹) to 1) was applied on each single cell spectrum.

#### Quantification of metabolic activities

Three metabolic activities were extracted from single-cell FTIR spectra. Protein synthesis rate was calculated as the ratio of the ¹³C-amino acid–labeled Amide I signal to the endogenous Amide I signal. Saturated fatty-acid metabolism was quantified from the azido-palmitic acid peak normalized to the total spectrum. Unsaturated fatty-acid metabolism was quantified from the deuterated oleic acid peaks normalized to the total spectrum.

### Optimal transport implementation and spectral pairing

#### Problem formulation

Each experimental condition as an empirical distribution over a common spectral space ***χ***, using:

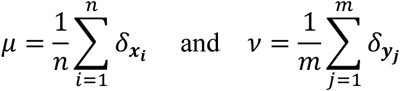

where *µ* and *v* represent the distributions of control and treated cell spectr respectively, with *x*_*i*_, *y*_j_ ∈ ***χ***. We hypothesized that drug treatment modifies only a subset of cellular metabolic programs, such that *v* can be approximated as a minimally shifted version of *µ* in spectral space.

#### OT coupling and perturbation vectors

To align *µ* and *v*, we adopted optimal transport (OT) as a principled approach for modeling perturbation-induced transformations of cell states. OT formalizes this alignment as an optimization problem that seeks a map *T*: ***χ*** → ***χ*** from control states to treated states, explicitly capturing the perturbation effect:

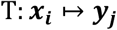

The classic Monge formulation seeks a deterministic mapping *T* that minimizes total spectral displacement when transporting mass from to *v*:

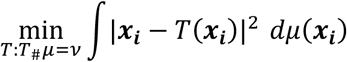

However, this deterministic formulation is often computationally intractable in discrete, high-dimensional settings. Thus, we adopted the Kantorovich relaxation, which allows soft, probabilistic assignments between the two distributions:

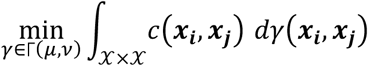

Here, *γ* in Γ(*µ*, *v*) denotes a coupling (or transport plan) between the empirical distributions *µ* and *v*, and *c*(***x*_*i*_**, ***x*_*j*_**) represents the transport cost, defined here as the squared Euclidean distance between spectral vectors.

The resulting OT solution is represented as a coupling matrix *γ* ∈ *R*^*n*×*m*^, where each entry *γ*_*i*j_ encodes the soft correspondence between control cell ***x*_*i*_** and treated cell ***y*_*j*_**. From this coupling, we computed cell-wise perturbation vectors:

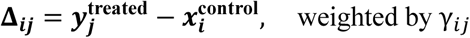

The resulting coupling forms the computational backbone of VIP-OT. It enables single-cell–level estimation of treatment-induced metabolic changes without requiring paired measurement.

#### OT solver implementation

The optimal transport problem was solved using the Python Optimal Transport (POT) library^90^. Cost matrices were computed directly in the original spectral space, without dimensionality reduction, to preserve the full biochemical information contained in vibrational profiles. Couplings were obtained as exact solutions to the Kantorovich formulation using the linear programming solver ot.emd, without entropic regularization. All OT analyses were performed in Python (v3.10) using POT (v0.9.2), with code and reproducibility scripts provided in the associated GitHub repository.

#### Analysis of baseline-dependent response heterogeneity

To stratify treated cells, we applied k-means clustering with k = 2 (scikit-learn) to obtain “low” and “high” response strata. Cluster labels were propagated from treated to control cells using the OT coupling. For each control cell, we normalized its outgoing OT weights to treated cells to sum to 1 and then assigned a single label by probability-proportional-to-weight sampling (one draw per source, fixed random seed). Baseline spectral features were then extracted on the control population as absorbance within predefined ROIs; Group differences (low vs high) were assessed with a two-sided Mann–Whitney U test. Effect size was summarized by Cliff’s delta. To evaluate predictive value, we computed ROC/AUC for classifying “low” vs “high” using baseline lipid ester alone (positive class = low; score = −ester), with stratified bootstrap 95% CIs (B = 5,000).

As diagnostics of the OT mapping, we examined (i) baseline vs barycentric expected post-treatment value and (ii) baseline vs change Δ (Δ = expected − baseline, summarizing concordance with Spearman’s ρ and Theil–Sen slopes with bootstrap CIs (Supplementary Fig. S5). These analyses verified rank preservation and the absence of baseline-dependent shifts under the OT pairing. All analyses were performed in Python (NumPy/Pandas/SciPy/Scikit-learn); random seeds were fixed at 0.

#### Single-cell drug response prediction

For each drug condition *d*, the model takes as input the baseline vibrational spectrum of an unperturbed control cell and outputs both the full post-treatment spectrum and three derived metabolic readouts: (1) protein synthesis rate; (2) saturated fatty acid metabolism; and (3) unsaturated fatty acid metabolism.

#### Training dataset generation

Given a control cell, *x*_*i*_ ∈ *R*^*d*^, denote the vibrational spectrum of the *i*-th control cell, and let 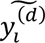 ∈ *R*^*d*^ represent its OT-weighted barycentric target under drug *d*. VIP-OT provides a soft coupling matrix 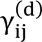 between control and treated populations. We constructed pseudo-targets for each control cell as OT-weighted treated averages:

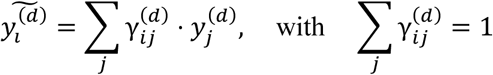

where 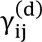 is the soft assignment weight from OT coupling, and 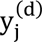 are treated cell spectra under drug *d*.

The resulting dataset for each drug consisted of pairs (*x*_*i*_, 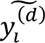), which served as training examples linking baseline spectra to estimated treatment outcomes. All drugs shared the same pool of control cells *X* = {*x*_1_, …, *x*_*n*_} that are shared across all drug conditions, ensuring a unified manifold across prediction tasks.

#### Prediction model

For each drug *d*, we sought a mapping *f*^(*d*)^: *x* → *ŷ*, that predicts the treated spectrum *ŷ* from baseline *x*. For each test control cell *x*_*test*_, we performed prediction via a nearest-neighbor strategy: we standardized all spectra and identified the training control cell *x*_train_ ∈ *X*_*train*_ that minimizes the Euclidean distance:

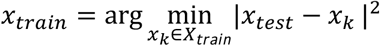

The predicted treated spectrum is then given by:

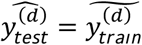

where 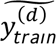 is the OT-weighted treated average associated with *x_train_* under drug *d*.

#### Cross-validation and evaluation

Each drug was processed independently under a 20-fold cross-validation scheme. In each fold, control cells were partitioned into training and test sets, and predictions were made for all held-out test cells. Prediction performance was quantified at two levels: (1) Spectrum-level accuracy: Pearson correlation between predicted and measured spectra, as well as R²; (2) Metabolic activities-level accuracy: R², mean absolute error (MAE), and mean squared error (MSE) for the three metabolic readouts derived from spectra.

For each drug, an overall prediction score was defined as the mean of spectrum-level R², spectral correlation, and average metabolic R².

#### Spectral Velocity Construction

To reconstruct dose-resolved trajectories from static vibrational spectra, we introduced the concept of Spectral Velocity, which estimates the most probable transition vector for each cell along a pseudo-temporal drug axis.

#### Embedding of spectral data

All single-cell spectra across a given dosage series were embedded into a shared low-dimensional latent space. In this work, we primarily used principal component analysis (PCA) for visualization and computational efficiency, though the framework is compatible with alternative embeddings (e.g., PHATE or UMAP). Each cell was thus represented by coordinates *z*_*i*_ ∈ *R*^*k*^, where *k* denotes the dimensionality of the embedding.

#### OT-derived displacement vectors

For each pair of adjacent dosage levels (e.g., *d*_1_ → *d*_2_), OT was applied to compute a soft coupling matrix γ_*i*j_ between cells at *d*_1_ and *d*_2_. For each source cell *x*_*i*_ at *d*_1_, its expected counterpart under *d*_2_was estimated as a barycentric projection:

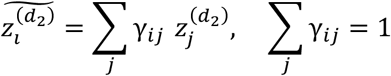

where 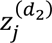 are the embedded coordinates of cells at dose *d*_2_.

#### Definition of spectral velocity

The spectral velocity for cell *i* was then defined as the weighted displacement vector:

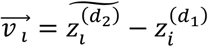

which represents the minimal displacement required for cell *i* to transition from its current state at dose *d*_1_ to its most probable future state at dose *d*_2_. For visualization, we restricted to high-confidence couplings (*max*_j_γ_*i*j_ ≥ 0.3) to reduce noise from uncertain pairings.

#### Velocity field visualization

Individual velocity vectors were plotted as arrows originating from the source cell positions in the latent space. To highlight coherent state transitions, we further interpolated these vectors using a kernel-based smoothing approach, generating streamline fields that depict global flow patterns across dosage levels.

#### Interpretation

The magnitude |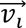| reflects the rate of spectral change, while the direction encodes the biochemical trajectory of the cell in latent space. By aggregating these vectors, we obtained pseudo-temporal maps of drug-induced metabolic remodeling at single-cell resolution.

## Acknowledgements

We thank the entire Min Lab and Shu Lab for their support and advice. This work was supported by NIH New Innovator Award (DP2TR004354, JS), NIH Common Fund Cellular Senescence Network (UH3CA275687, UG3CA275687, JS), NIH (P01AI129859, JS), Massachusetts Life Science Center (JS), Burroughs Wellcome Fund (JS), Additional Ventures (JS), and Massachusetts General Hospital (JS). W.M. acknowledges support from the National Institute of Health (R35 GM149256) and Chan Zuckerberg Initiative (Dynamic Imaging 2023-321166).

## Contributions

JS and WM conceived, designed, and supervised the study. XML, SGS, XWL designed the experiments. XML, MW performed the experiments. XML, MW processed the single-cell spectra. XML, SGS performed the data analysis. XML, WM, JS wrote the paper with input from all authors.

## Code Availability

The codes are available at https://github.com/jian-shu-lab/VIP-OT.

## Declaration of interests

A patent application based on this study has been filed. All authors declare no conflict of interest.

## Supplementary Information

**Supplementary Figure 1.**
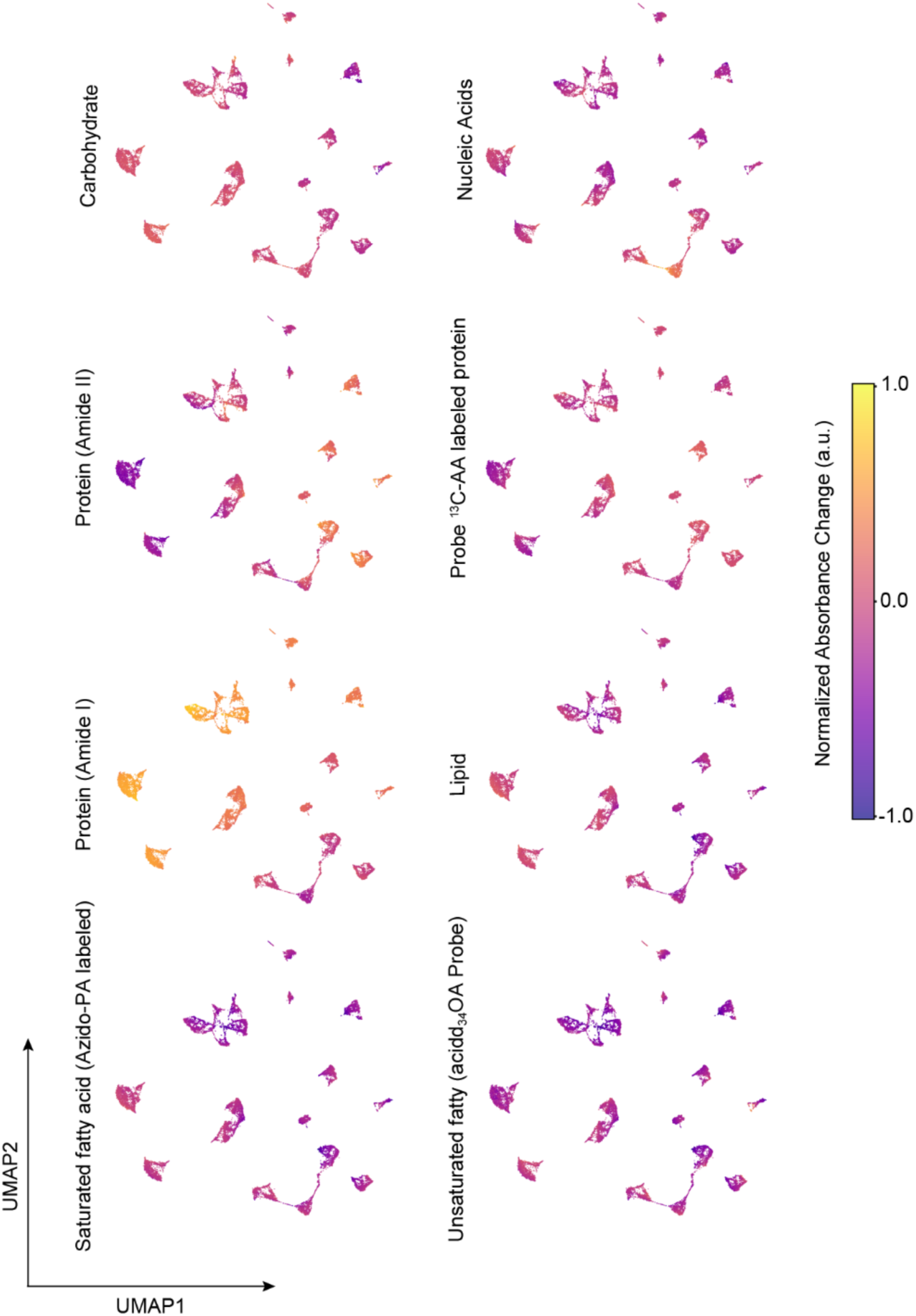
UMAP projection of treated cells colored by absorbance within selected spectral bands associated with major molecular classes. Spatial distribution of probe signals reveals MoA-consistent suppression or enhancement in protein and lipid-related regions.

**Supplementary Figure 2.**
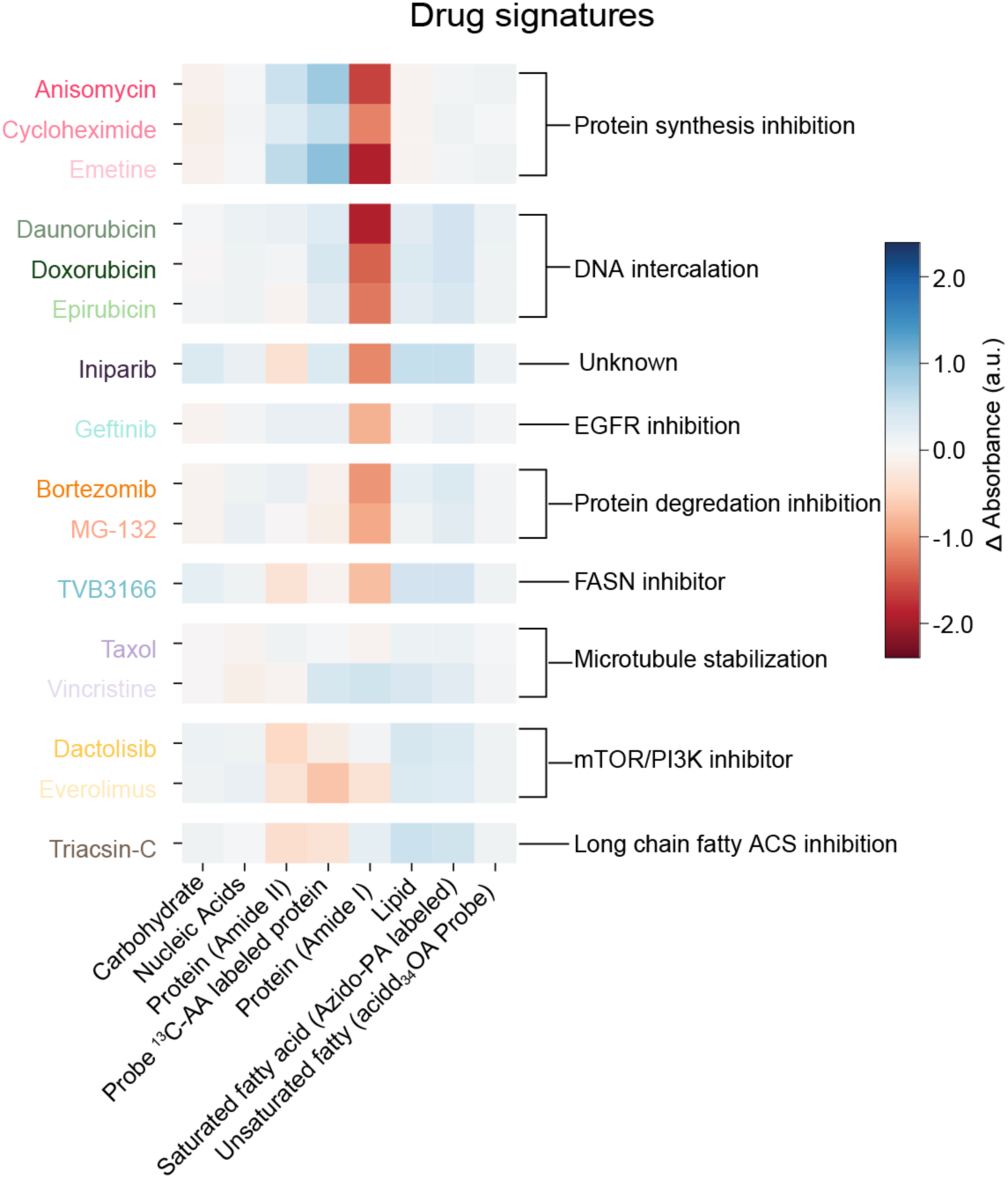
Heatmap of drug signatures derived by averaging difference spectra across probe-sensitive and endogenous spectral bands. Each row corresponds to a drug, and each column denotes a spectral region, capturing distinct biochemical fingerprints associated with specific perturbations.

**Supplementary Figure 3.**
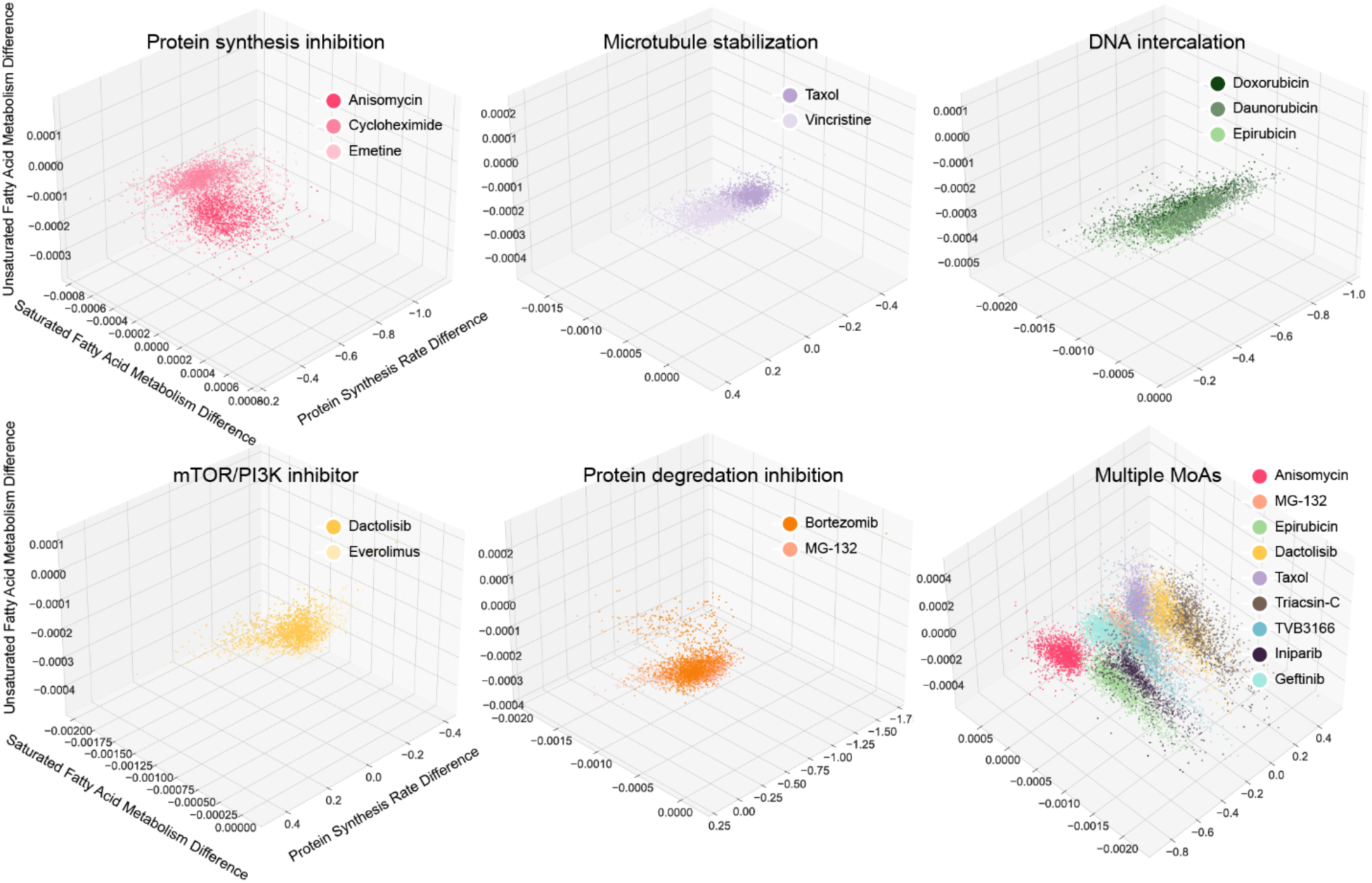
Metabolic coordinate plots of drug-induced perturbation effects, with each point representing a paired cell’s difference in protein synthesis, saturated fatty acid metabolism, and unsaturated fatty acid metabolism.

**Supplementary Figure 4.**
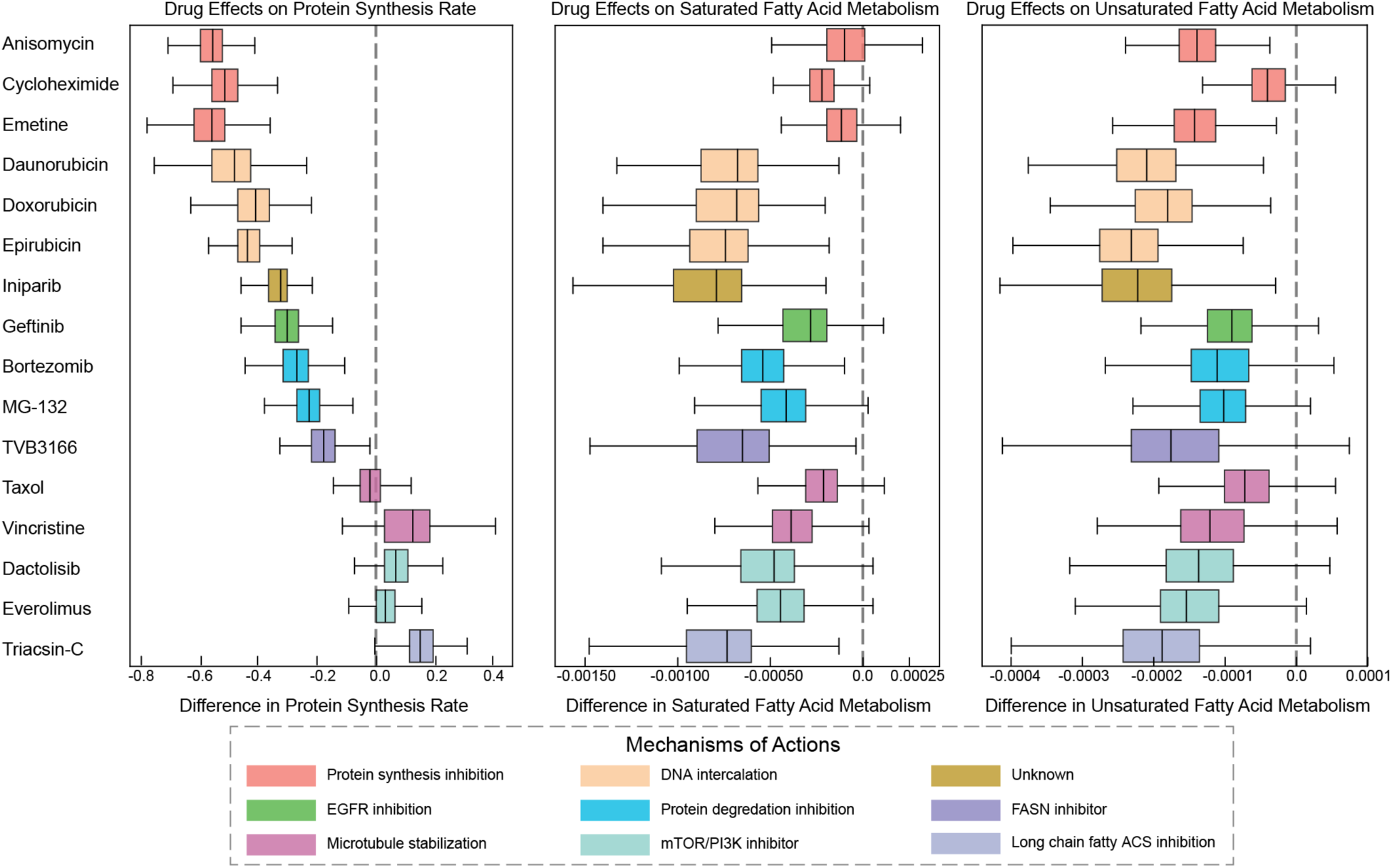
Boxplots showing the per-cell distribution of metabolic changes across three axes, protein synthesis, saturated fatty acid metabolism, and unsaturated fatty acid metabolism, for all drugs. Each box reflects the effect size derived from VIP-OT pairing, with colors indicating MoA groups and dashed lines marking zero-change baselines.

**Supplementary Figure 5.**
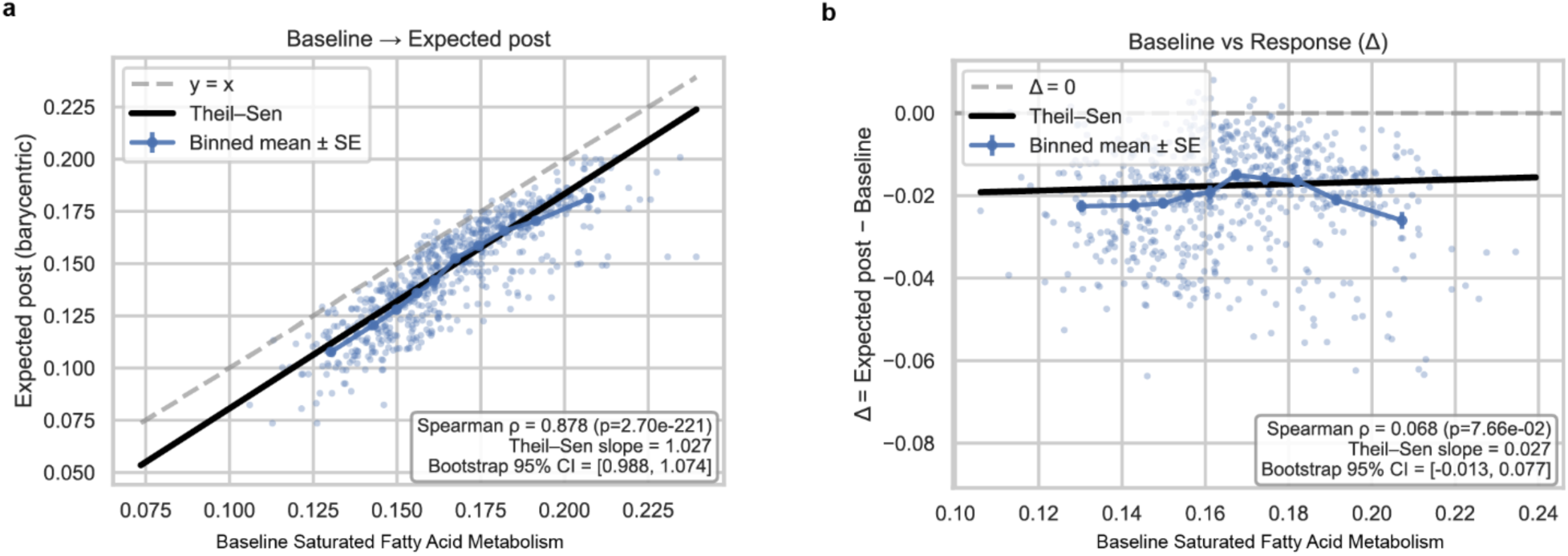
Barycentric mapping diagnostics. **a,** Scatter of baseline vs barycentric expected post-treatment metric shows a strong monotonic relation (Spearman ρ = 0.878) and a trend slope ≈ 1.03 (95% CI 0.988–1.074), close to the identity line, which is consistent with rank preservation and near-proportional scaling. **b,** Scatter of baseline vs response Δ (Δ= expected − baseline) shows no association (ρ = 0.068) and a near-zero slope (0.027; 95% CI −0.013–0.077), indicating the change is not driven by baseline. Points represent control cells; per-source OT weights sum to 1.

**Supplementary Figure 6.**
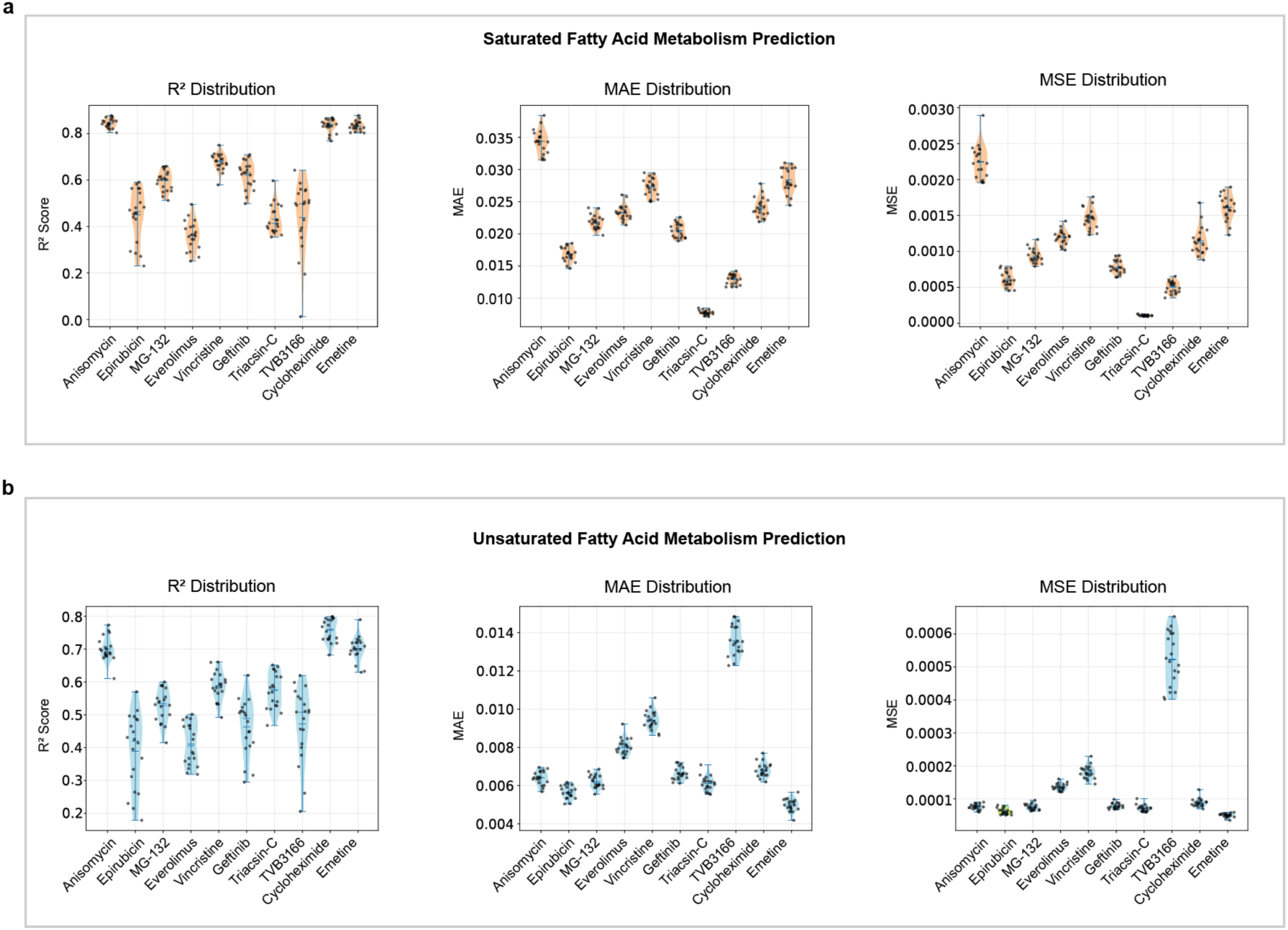
Violin plots showing R², MAE, and MSE distributions of prediction performance for saturated fatty-acid metabolism (a) and unsaturated fatty-acid metabolism (b) across 10 drugs and 20-fold cross-validation. Results are shown in analogy to Fig. 4b, where protein synthesis rate predictions are presented.

**Supplementary Figure 7.**
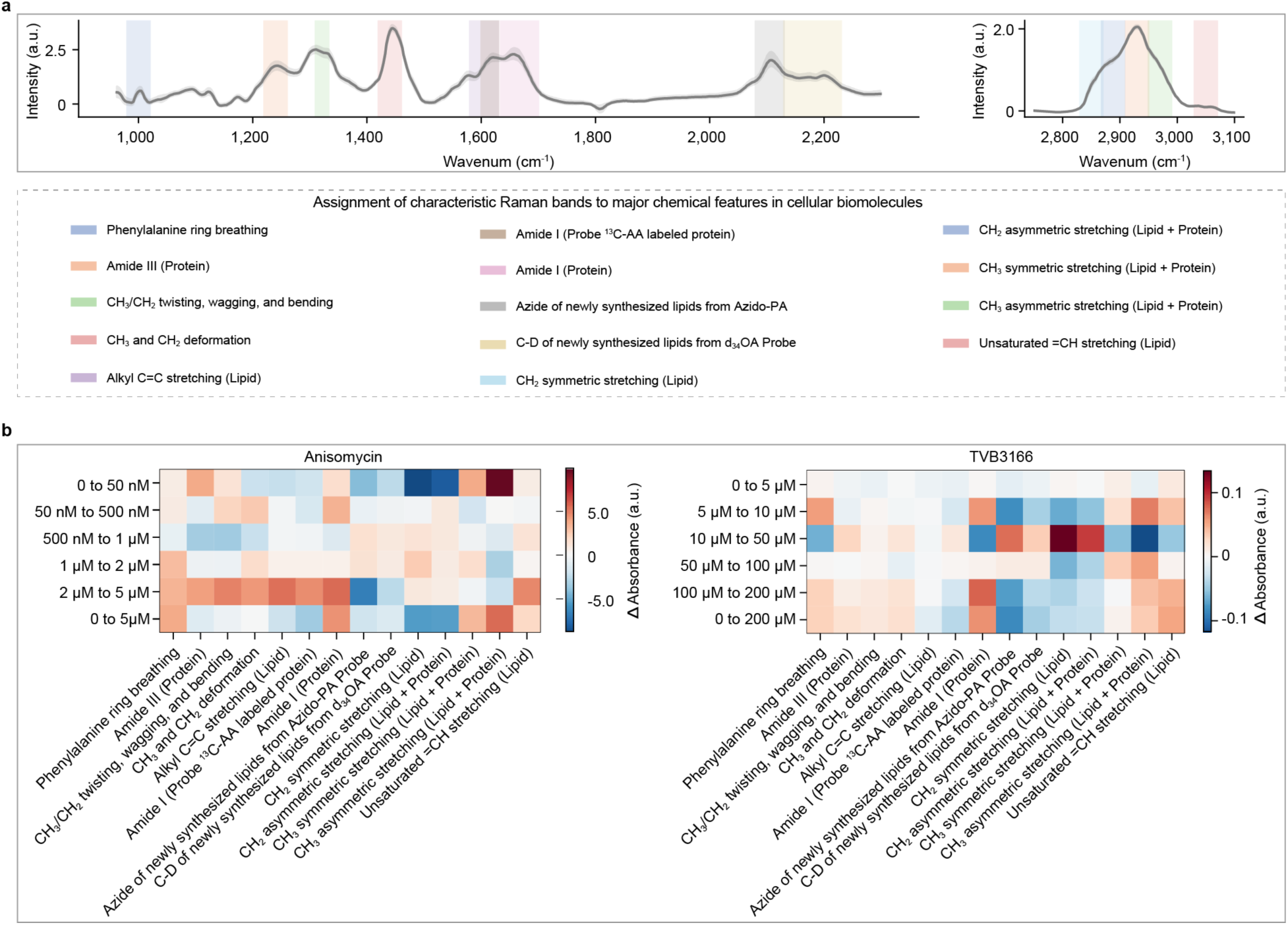
Raman measurements of drug-treated MDA-MB-231 cells. **a,** Mean spontaneous Raman spectra of DMSO control cells with shaded regions indicating 95% confidence intervals, plotted separately for the fingerprint (900–2300 cm⁻¹, left) and CH-stretching (2700–3100 cm⁻¹, right) regions due to differences in signal intensity. Colored bands highlight vibrational modes corresponding to major molecular classes. **b,** Heatmaps summarizing Raman-derived spectral signatures across dosage intervals for anisomycin and TVB-3166.

**Supplementary Fig 8.**
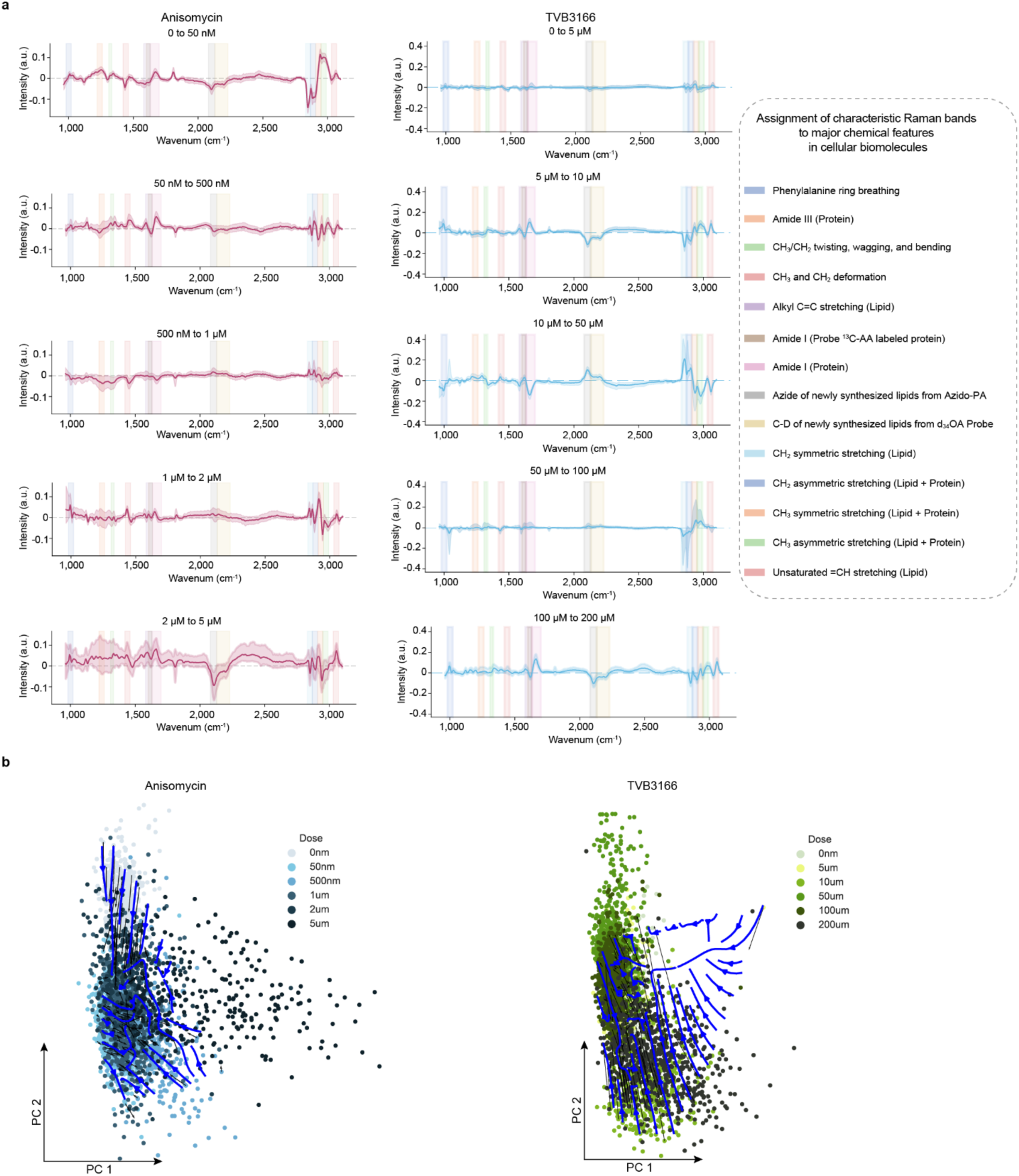
Raman-based validation of VIP-OT trajectory modeling. a, Raman difference feature spectra between adjacent dose levels for anisomycin and TVB-3166. **b,** Spectral Velocity Maps constructed from OT-derived displacement vectors in a PCA embedding. Black arrows represent sampled displacement vectors from high-confidence OT pairings (≥ 0.3), and blue streamlines indicate interpolated velocity fields.

**Supplementary Table 1.**
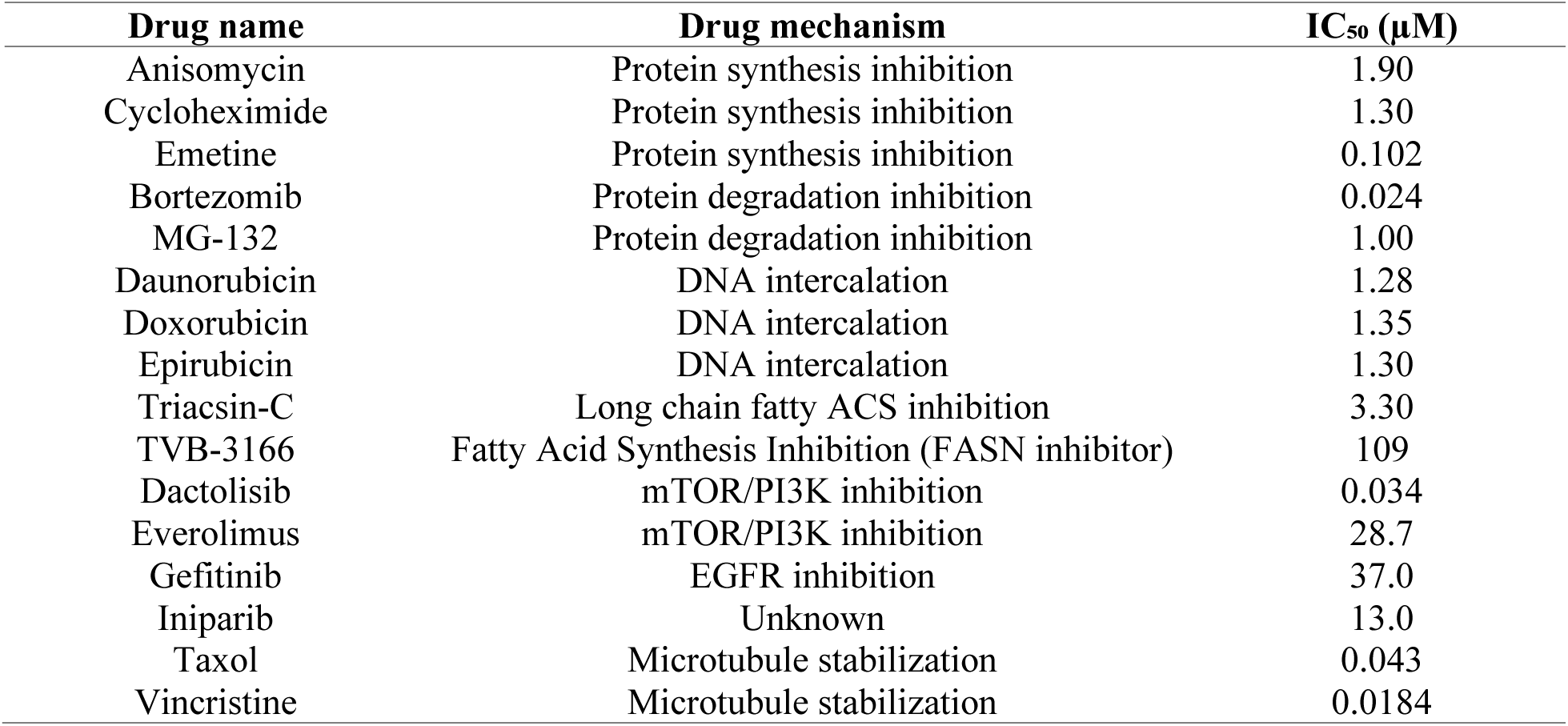
List of 16 drugs used in the VIP-OT validation dataset. Drugs are grouped into two main categories: inhibitors of general metabolism (e.g., protein synthesis, DNA intercalation, proteasome inhibition) and agents targeting specific signaling pathways (e.g., PI3K/mTOR, EGFR, PARP). For each drug, the mechanism of action (MoA), target pathway, and concentration applied (IC₅₀) are indicated.

| Drug name | Drug mechanism | IC<sub>50</sub> (μM) |
| --- | --- | --- |
| Anisomycin | Protein synthesis inhibition | 1.90 |
| Cycloheximide | Protein synthesis inhibition | 1.30 |
| Emetine | Protein synthesis inhibition | 0.102 |
| Bortezomib | Protein degradation inhibition | 0.024 |
| MG-132 | Protein degradation inhibition | 1.00 |
| Daunorubicin | DNA intercalation | 1.28 |
| Doxorubicin | DNA intercalation | 1.35 |
| Epirubicin | DNA intercalation | 1.30 |
| Triacsin-C | Long chain fatty ACS inhibition | 3.30 |
| TVB-3166 | Fatty Acid Synthesis Inhibition (FASN inhibitor) | 109 |
| Dactolisib | mTOR/PI3K inhibition | 0.034 |
| Everolimus | mTOR/PI3K inhibition | 28.7 |
| Gefitinib | EGFR inhibition | 37.0 |
| Iniparib | Unknown | 13.0 |
| Taxol | Microtubule stabilization | 0.043 |
| Vincristine | Microtubule stabilization | 0.0184 |

**Supplementary Table 2.** Drug concentrations used for VIP-OT dose–response FTIR experiments. For each drug, MDA-MB-231 cells were treated at the indicated concentrations; details of experimental setup are provided in Methods.

| Drug | MoA | Concentrations used (μM) |
| --- | --- | --- |
| Anisomycin | Protein Synthesis Inhibition | 0.05, 1.0, 5.0 |
| TVB-3166 | Fatty Acid Synthesis Inhibition | 5, 10, 200 |

**Supplementary Table 3.**
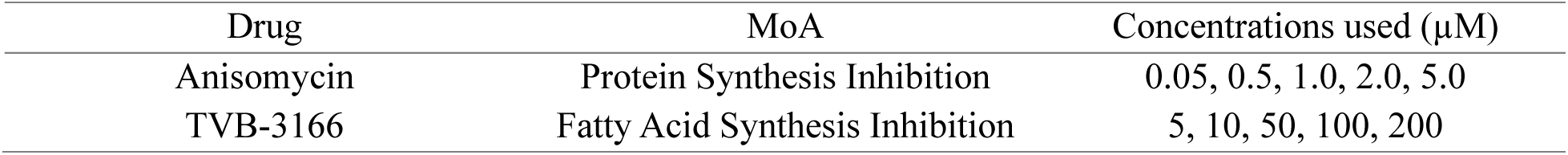
Drug concentrations used for VIP-OT dose–response spontaneous Raman experiments. For each drug, MDA-MB-231 cells were treated at the indicated concentrations; details of experimental setup are provided in Methods.

**Supplementary Table 4.** Concentrations of Bortezomib (Bor) and Gefitinib (Gef) used in VIP-OT drug combination experiments. Conditions included single-agent treatments at IC_20_ and IC_50_ as well as matched combinations, as described in Methods.

| Condition | Bortezomib conc. (μM) | Gefitinib conc. (μM) | Notes |
| --- | --- | --- | --- |
| Control | 0 | 0 | Untreated |
| Bor IC<sub>20</sub> | 0.0130 | 0 | Monotherapy |
| Bor IC<sub>50</sub> | 0.0240 | 0 | Monotherapy |
| Gef IC<sub>20</sub> | 0 | 24.0 | Monotherapy |
| Gef IC<sub>50</sub> | 0 | 37.0 | Monotherapy |
| Combo IC<sub>20</sub>/IC<sub>20</sub> | 0.0130 | 24.0 | Combination |
| Combo IC<sub>20</sub>/IC<sub>50</sub> | 0.0130 | 37.0 | Combination |
| Combo IC<sub>50</sub>/IC<sub>20</sub> | 0.0240 | 24.0 | Combination |
| Combo IC<sub>50</sub>/IC<sub>50</sub> | 0.0240 | 37.0 | Combination |

